# The establishment of multiple knockout mutants of *Colletotrichum orbiculare* by CRISPR/Cas9 and Cre/*loxP* systems

**DOI:** 10.1101/2021.10.24.465644

**Authors:** Kohji Yamada, Toya Yamamoto, Kanon Uwasa, Keishi Osakabe, Yoshitaka Takano

## Abstract

Phytopathogenic fungi belonging to the *Colletotrichum* genus cause devastating damage for many plant species. Among them, *Colletotrichum orbiculare* is employed as a model fungus to analyze molecular aspects of plant-fungus interactions. Although gene disruption via homologous recombination (HR) was established for *C. orbiculare*, this approach is laborious due to its low efficiency. Here we developed methods to efficiently generate multiple knockout mutants of *C. orbiculare*. We first found that CRISPR/Cas9 system massively promoted gene-targeting efficiency. By transiently introducing a CRISPR/Cas9 vector, more than 90 % of obtained transformants were knockout mutants. Furthermore, we optimized a self-excision Cre/*loxP* marker recycling system for *C. orbiculare* because limited availability of desired selective markers hampers sequential gene disruption. In this system, integrated selective marker is removable from the genome via Cre recombinase driven by a xylose-inducible promoter, enabling reuse of the same selective marker for the next transformation. Using our CRISPR/Cas9 and Cre/*loxP* systems, we attempted to identify functional sugar transporters in *C. orbiculare*. Multiple disruptions of putative quinate transporter genes restrict fungal growth on media containing quinate as a sole carbon source, confirming their functionality as quinate transporters. Our analyses revealed that quinate acquisition is dispensable during fungal infection because this mutant displayed normal virulence to host plants. In addition, we successfully built mutations of 17 cellobiose transporter genes in a strain. From the data of knockout mutants established in this study, we inferred that repetitive rounds of gene disruption using CRISPR/Cas9 and Cre/*loxP* systems do not cause negative effects for fungal virulence and growth. Therefore, these systems will be powerful tools to perform systematic gene targeting approach for *C. orbiculare*.

## Introduction

The ascomycete genus *Colletotrichum* represents one of economically important phytopathogenic fungal groups that infect a wide range of plants including commercial crops. Due to their worldwide occurrence, *Colletotrichum* species are ranked in top 10 important phytopathogenic fungi (*1*). Of them, *Colletotrichum orbiculare*, the causal agent of anthracnose on cucurbits, has been employed as a model fungus to analyze molecular aspects of plant-fungus interactions (*2*). In molecular biology, functional analysis of a particular gene often relies on gene manipulation experiments, especially by gene disruption. Methods of homologous recombination (HR)-based gene replacement were developed for *C. orbiculare* although these classical procedures are laborious due to its low efficiency. The attempts to increase HR efficiency in filamentous fungi were reported. Exogenous DNA is thought to be integrated into a chromosome via DNA repair systems (*3*). Two distinct pathways are known to repair double-strand breaks (DSBs): NHEJ pathway which aligns and ligates broken DNA ends without long homologous sequences and HR pathway which requires sequences homologous with the broken DNA as a template. Because these two pathways are considered to act independently and competitively, disruption of NHEJ pathway promoted HR efficiency in many fungi (*4–9*). On the other hand, DSBs on targeted loci rarely occur. Therefore, artificial DSB introduction using genome editing technologies have been applied to promote HR efficiency in fungi. Of genome editing tools, the clustered regularly interspaced short palindoromic repeats (CRISPR) –associated RNA-guided Cas9 endonuclease becomes the leading tool to recruit nucleases on specific loci to introduce DSBs in various organisms(*10–12*). Elevation of HR efficiency using CRISPR/Cas9 system was reported in filamentous fungi including phytopathogenic fungi such as *Magnaporte oryzae* (*13*) and *Botrytis cinerea* (*14*).

In recent years, the genomes of filamentous fungi including *Colletotrichum* species have been sequenced (*15, 16*) These studies revealed the presence of an excess of homologous genes, indicating high degrees of their functional redundancy. Therefore, genes belong to such gene families need to be disrupted on comprehensive scales to address their cellular functions. However, a limited number of available selective markers hampers sequential transformations. To circumvent this problem, marker recycling systems, in which introduced marker genes are removed from the genome, have been developed. Two recombination systems are primarily used in marker recycling systems, the bacteriophage P1-derived Cre/*loxP* system (*17*) and the *Saccharomyces cerevisiae FLP/FRT* system (*18*). Tyrosine recombinase Cre and FLP binds to *loxP* and *FRT* sequence, respectively, to catalyze recombination, resulting in excision, insertion, translocation and inversion of DNA fragments. By optimizing these systems, a marker gene located within *loxP* and *FRT* can be removed by Cre and FLP, respectively. These approaches were also applied to phytopathogenic fungi such as *Fusarium graminearum* (*19*) and *Ustilago maydis* (*20*).

In this study, we engineered CRISPR/Cas9 system and Cre/*loxP* system to enhance HR-based gene disruption and to generate multiple knockout (KO) strains, respectively, for *C. orbiculare*. Especially, we developed a self-excision Cre/*loxP* system to reduce laborious experimental steps to generate multiple KO strains. In this system, *Cre* gene fragment is located together with positive/negative selective markers within two *loxP* sequences, and this marker cassette can be removed by Cre driven by xylose-inducible promoter. By using these tools, we identified quinate transporters in *C. orbiculare*. Furthermore, we successfully established mutants lacking 17 cellobiose transporter genes. Altogether, we believe that these systems will be helpful to perform systematic gene targeting approach for *C. orbiculare*.

## Results

### CRISPR/Cas9 system promotes gene-targeting efficiency in *C. orbiculare*

Although HR-based gene targeting has been applied to *C. orbiculare*, its efficiency is low. In this study, we first tried to promote efficiency of gene targeting by introducing DSBs on the targeted locus via CRISPR/Cas9 system. To easily evaluate gene-targeting efficiency, *PKS1* gene which encodes polyketide synthase (*21, 22*) was disrupted. Because *pks1* strain becomes orange color phenotype due to a lack of melanin biosynthesis, we can distinguish *pks1* mutants from the wild-type (WT) strain without PCR-based genotyping (Fig. 1A). For gene targeting of *PKS1* gene, we amplified 1 kbp fragments of 5’- and 3’-flanking sequences of *PKS1* coding region, and fused *hygromycin phosphotransferase* (*HPT*) gene (hygromycin resistance gene) driven by the constitutive *TrpC* promoter (pCB1636 *pks1*) (Fig. 1B). For introducing CRISPR/Cas9 system (pChPtef026 vector), codon-optimized *Cas9* sequence and gRNA is driven by *Aureobasidium pullulans translation elongation factor 1-alpha* (*tef*) constitutive promoter and *Colletotrichum higginsianum U6* promoter, respectively (Fig. 1B and Fig. S2). Cas9 was C-terminally fused with three-repeated nuclear localization signals. We designed gRNA sequences from *PKS1* locus and ligated to pChPtef026 vector (pChPtef026 *pks1*) (Fig. 1B and 1C). Disruption of NHEJ pathway, for example by mutating *Lig4* or *Ku70*, was reported to promote HR efficiency in many fungi (*4–9*). Although the *lig4* strain of *C. orbiculare* was recently generated and utilized for the gene knockout analysis, (*23*), HR efficiency of this strain has not been evaluated yet. Therefore, we co-transformed two plasmids, targeting vector (pCB1636 *pks1*) and CRISPR/Cas9 vector (pChPtef026 *pks1*), to WT strain (104-T) and *lig4* strain. Without *PKS1* gRNA in pChPtef026 vector, only one out of five transformants were *pks1* mutants in WT strain (Fig. 1D). On the other hands, three out of seven transformants were *pks1* mutants in *lig4* strain (Fig. 1D). This result suggested that loss of *LIG4* gene improved gene targeting efficiency in *C. orbiculare*. However, this efficiency was lower than expected because disruption of *Ku70* led to much higher HR efficiency in *C. higginsianum* (*4*). We next used pChPtef026 vector containing *PKS1* gRNA1 (5’-GTAGGTGTCGACCTCTTGAG-3’). Surprisingly, the number of transformants markedly increased, and most of them (more than 90 %) were *pks1* mutants in both WT and *lig4* strains (Fig. 1D). On the other hand, the effect of gRNA2 (5’-GATGTCGTAGTGGGTAGCGA-3’) on gene targeting efficiency was limited, compared to gRNA1, suggesting that gRNA sequence largely influences HR efficiency. In addition, we found that the KO rate using gRNA2 was higher in *lig4* stain than WT stain, also indicating that disruption of NHEJ pathway facilitates HR-based gene replacement in *C. orbiculare*.

**Fig. 1.**
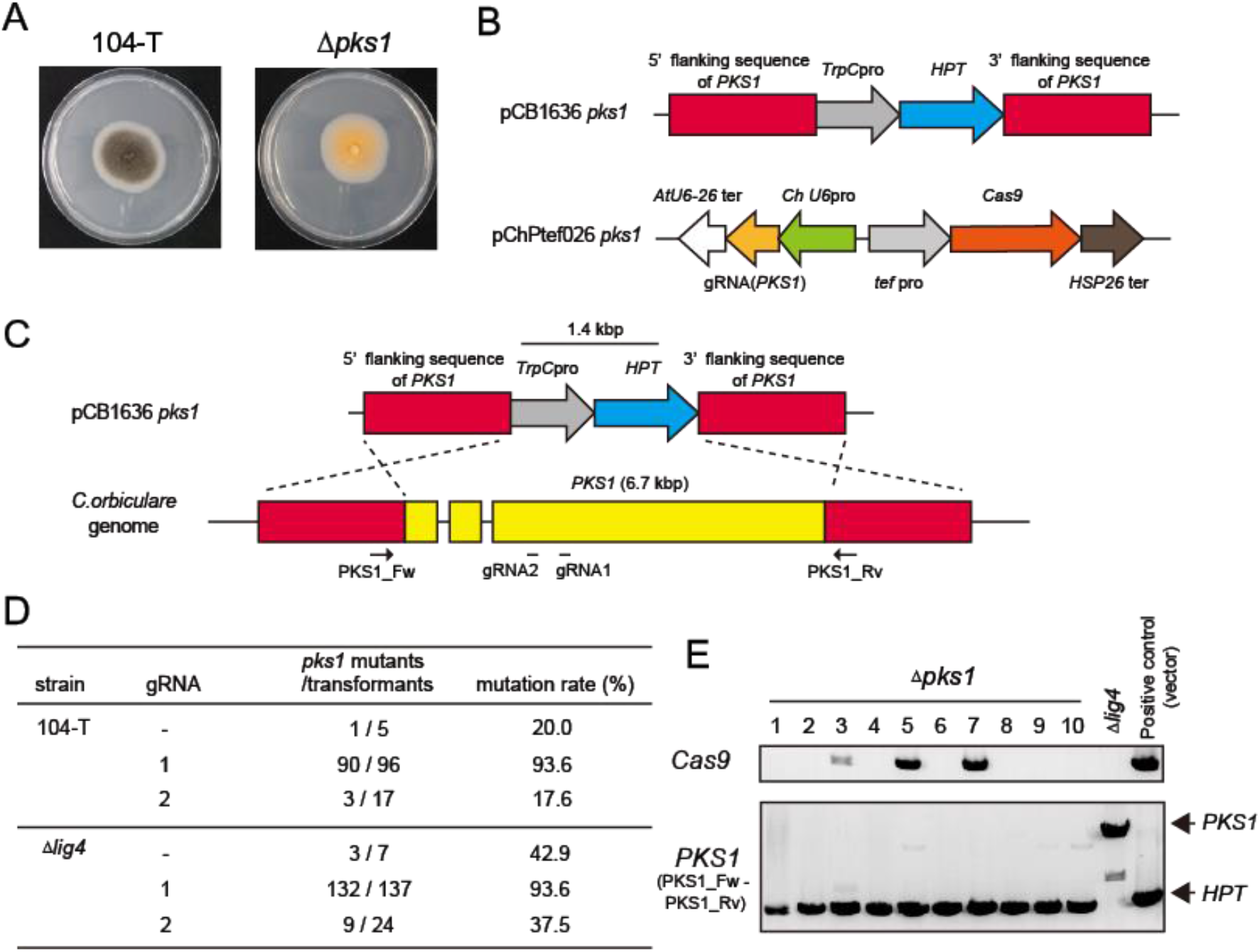
CRISPR/Cas9 system promotes gene-targeting efficiency in *C. orbiculare*. A, Phenotype of *pks1* mutants. *pks1* mutants show orange color phenotype due to lack of melanin synthesis. B, Vector information of the targeting vector pCB1636 *pks1* and the CRISPR/Cas9 vector pChPtef026 *pks1*. C, Scheme of gene disruption using the targeting vector pCB1636 *pks1*. The positions of gRNA were lined under *PKS1* locus. D, Efficiency of establishment of *pks1* mutants using CRISPR/Cas9 system. Results of three independent experiments were combined. E, Insertion of *Cas9* fragment in the genome of *pks1* mutants generated using pCB1636 *pks1* and pChPtef026 *pks1*.

Because pChPtef026 vector does not have any selective markers for *C. orbiculare*, we next investigated if *Cas9* is integrated into the genome or transiently expressed during transformation. *Cas9* fragments was detected in three out of ten *pks1* mutants (Fig. 1E), indicating that transient *Cas9* expression is basically sufficient to promote gene-targeting efficiency. Genome integration of CRISPR/Cas9 system hampers complementation assay for KO strains. Therefore, we concluded that this system can be a strong tool for loss-of-function study in in *C. orbiculare*.

### Cre/*loxP* system works in *C. orbiculare*

Proteins which share a common evolutionary origin shape groups known as protein families. Due to their redundant functions, multiple genes in a protein family often need to be disrupted for loss-of-function study. However, a limited number of available selective markers restricts sequential transformation in *C. orbiculare*. Previous studies reported that recyclable marker modules allow repetitive rounds of gene disruption in fungi (*20, 24*). Therefore, we investigated whether Cre/*loxP* system-based marker recycling is applicable to *C. orbiculare*. Cre recombinase evicts a DNA fragment within two 34 bp *loxP* sequences. We constructed the plasmid (pCB1636 lox HPT-TK) in which the fragment of *HPT-TK* gene, the positive marker *HPT* gene fused with the negative maker *thymidine kinase (TK*) gene, driven by *TrpC* promoter is sandwiched with *loxP* sequences (Fig. 2A). In Cre/*loxP* system, single *loxP* sequence is remained in the genome after Cre-based recombination, possibly causing unfavorable arrangement in the next rounds to build multiple mutations. Therefore, we employed mutated *loxP* sequences, *lox66* and *lox77* (*25*). By using these *loxP* sequences, remained *loxP* sequence becomes inefficient *lox72* sequence. We transformed pCB1636 lox HPT-TK into WT protoplasts and selected transformants with hygromycin. Sequentially, we transformed pChPtef026 or pChPtef026 Cre vector in which codon-optimized *Cre* gene is driven by the *tef* promoter, and incubated transformants with 5-fluoro-2’-deoxyuridine (FdU). Because TK converts FdU to be a toxic compound, only colonies which do not have *TK* gene are able to grow on PDA media containing FdU. No colonies were detectable after transforming the vector without *Cre* gene (pChPtef026), indicating TK-mediated negative selection works in *C. orbicuolare* (Fig. 2B). On the other hand, colonies became observable after the introduction of pChPtef026 Cre vector (Fig. 2B). We confirmed the removal of *HPT-TK* fragment using PCR (Fig. 2C). Importantly, Cre-introduced strains became sensitive to hygromycin and insensitive to FdU like the WT strain (Fig. 2D).

**Fig. 2.**
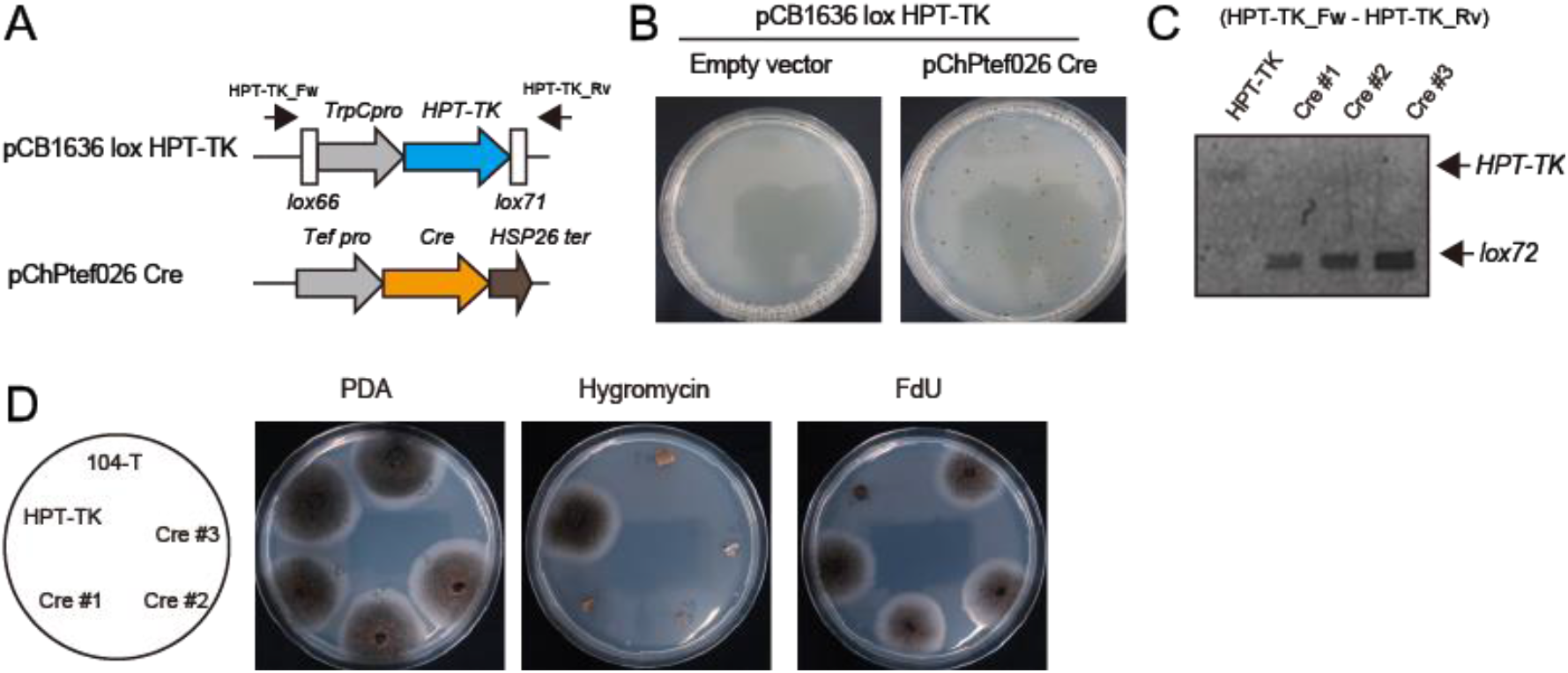
Cre/*loxP* system works in *C. orbiculare*. A, Vector information of pCB1636 lox HPT-TK and pChPtef026 Cre. B, The introduction of *thymidine kinase* (*TK*) gene led to inhibition of fungal growth on PDA containing FdU (left). Fungal colonies became observable after Cre introduction (right). C, Cre removed the DNA fragments containing *HPT-TK* gene within *loxP* sequences. D, Hygromycin resistance was lost in Cre-introduced strains.

### Optimization of self-excision Cre/*loxP* systems for *C. orbiculare*

These results indicated that Cre/*loxP*-based marker recycling is applicable to *C. orbiculare*. However, this system is laborious and time-consuming because two transformation steps, gene targeting and Cre introduction, are required to disrupt one gene. We next tried to optimize a self-excision Cre/*loxP* system for *C. orbiculare*. In this system, *Cre* gene is also inserted within *loxP* sequences, and Cre protein removes *Cre* gene itself together with *HPT-TK* gene (Fig. 3A). For this system, an inducible promoter to control *Cre* expression is required although no inducible promoters were reported in *C. orbiculare*. Because xylose-inducible promoters were popularly used for marker recycling systems in fungi (*18, 26–28*), we investigated xylose-inducibility of previously published *Penicillium chrysogenum xylP* promoter and *Aspergillus oryzae xynG2* promoter by *GFP* expression in *C. orbiculare*. Although both *PcxylP* promoter and *AoxynG2* promoter showed strong GFP induction on minimal media containing 2 % xylose media as a sole carbon source, GFP fluorescence was detectable even on PDA media without xylose (Fig. 3B). Because leaky *Cre* expression may cause a removal of the selective cassette in an inappropriate timing, we attempted to find other promoters which display weaker basal expression. We searched for xylase genes in *C. orbicualre*, and found two genes *Cob_01864* and *Cob_02882* which are named *xyl1* and *xyl2*, respectively. In *Neurospora crassa*, the expression of xylase gene is regulated by the transcription activator XLR-1 (*29*). We found putative XLR-1 binding sites, GGCTRR and GGNTAAAA (*29, 30*), in both promoters (Fig. S1), suggesting their xylose inducibility. We amplified 2 kbp upstream sequences from their coding regions as their promoters. Putative XLR-1 binding sites were found more in *Coxyl1* promoter than *Coxyl2* promoter in this region (Fig. S1). *Coxyl1* promoter expectedly showed stronger GFP expression on xylose media than *Coxyl2* promoter (Fig. 3B). Importantly, both promoters showed very low basal GFP expression without xylose (Fig. 3B). We employed *Coxyl1* promoter to regulate *Cre* gene. However, we failed to construct vectors for self-excision Cre/*loxP* systems likely due to leaky *Cre* expression in *E.coli*. To circumvent this problem, an intron-containing *Cre* gene was previously used (*31*). We here inserted an intron of *C. orbiculare histon h4* gene to *Cre* gene, and named it *Cre_i*. We confirmed that Cre_i worked as well as Cre in *C. orbiculare* (Fig. 3C).

**Fig. 3.**
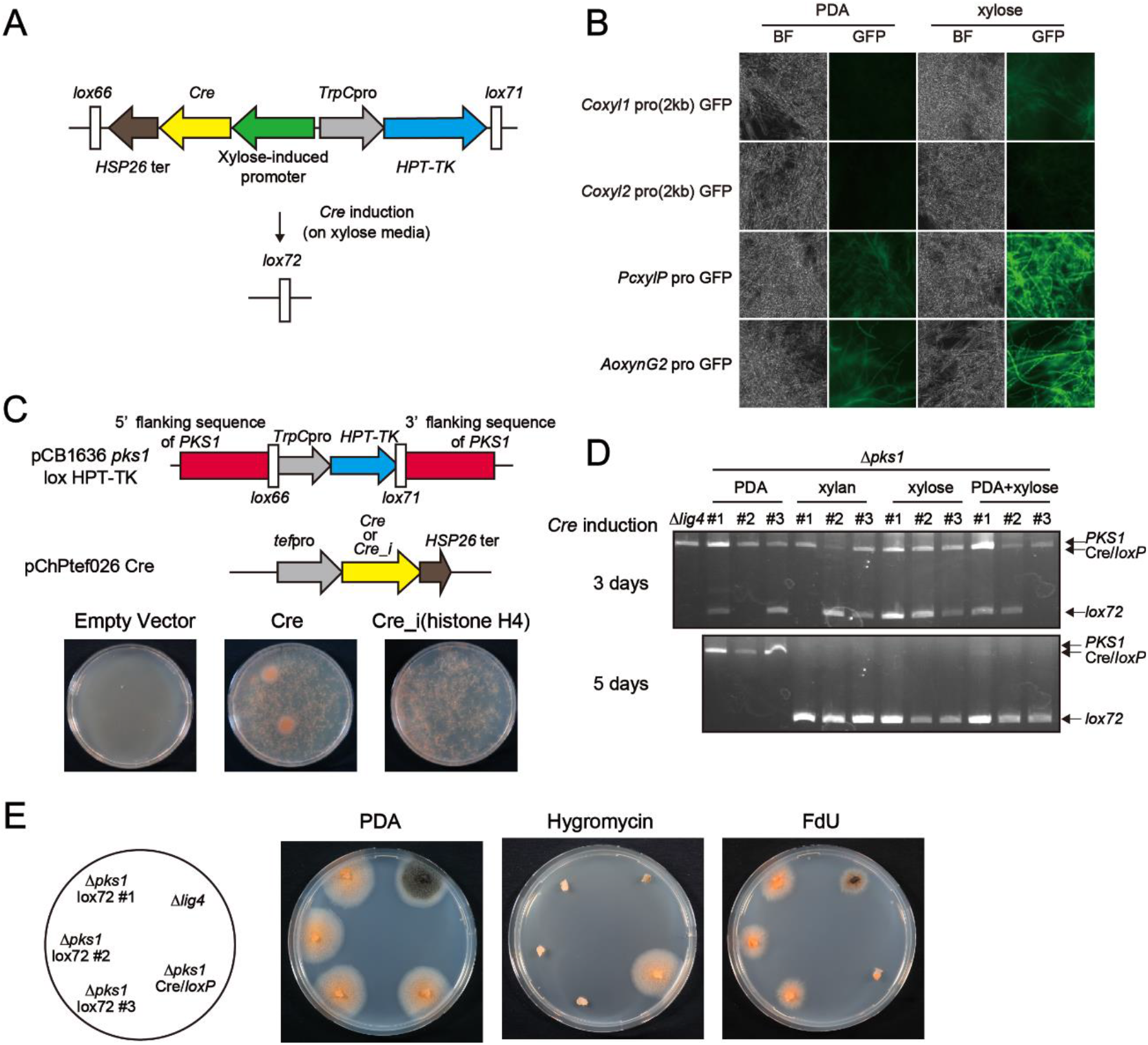
Establishment of a self-excision Cre/*loxP* system for *C. orbiculare*. A, Scheme of a self-excision Cre/*loxP* system in this study. Cre gene is induced by a xylose-inducible promoter, leading to excision of the DNA fragment within *loxP* sites. B, Xylose-inducibility of xylase gene promoters in *C. orbiculare*. BF indicates bright field. C, The insertion of an intron did not affect functionality of Cre in *C. orbiculare* D, Cre/*loxP* system is activated on media containing xylose or xylan. Cre removed Cre/*loxP* fragment, leading to generation of *lox72* fragment. E, Selective-marker cassette was successfully removed by Cre/*loxP* system.

We constructed a self-excision Cre/*loxP* plasmid to generate *pks1* mutants. After establishing *pks1* mutants, these *pks1* mutants were grown on minimal media containing xylan or xylose (Fig. 3D). Because xylase plays a role to hydrolyze xylan, *Coxyl1* promoter might become more active in the presence of xylan than xylose. However, *lox72* fragment which is indicative of successful Cre-based eviction was detectable to a similar extent on both xylan and xylose media (Fig. 3D). Because the presence of glucose leads to inhibition of catabolite processes of other sugars, known as catabolite repression, we next tested if xylose effect was inhibited on PDA media. Unexpectedly we found that Cre/*loxP* system worked on PDA media containing xylose (PDA+xylose) (Fig. 3D). The presence of other sugars did not restrict xylose inducibility of *Coxyl1* promoter. Therefore, minimal media does not need to be used for Cre induction. On the other hand, *lox72* fragments were also detectable on PDA media without xylose or xylan in some cases (Fig. 3D), indicating that the basal activity of *Coxyl1* promoter was weak but not completely off. Cre/*loxP* fragments was markedly reduced after 5 days on inducible media. Therefore, we decided to treat fungal strains on PDA+xylose media for 5 days to remove Cre/*loxP* cassette.

The conidia, harvested from *pks1* mutants after grown for 5 days on PDA+xylose media, were streaked on PDA containing FdU to pick up single colonies which do not have the Cre/*loxP* cassette. We named this strain *pks1* lox72. Although these strains showed orange color *pks1* phenotype, they lost hygromycin resistance (Fig. 3E). These results indicates that self-excision Cre/*loxP* system for *C. orbicualre* was successfully established.

We next tried to improve this system towards being more easily handled and efficient for establishing multiple mutants. Because this self-excision Cre/*loxP* fragment is a bit too long to be handled for plasmid construction, we attempted to reduce this fragment size. We investigated if a shorten *Coxyl1* promoter (650 bp) containing several putative XLR-1 binding sites was also functional for Cre induction on PDA+xylose media. Eviction efficiency seemed to be slightly reduced in the 650 bp *Coxyl1* promoter compared to 2kbp one (Fig. 4A). However, because Cre/*loxP* system surely worked, we employed this shorten version of *Coxyl1* promoter for our self-excision Cre/*loxP* system.

**Fig. 4.**
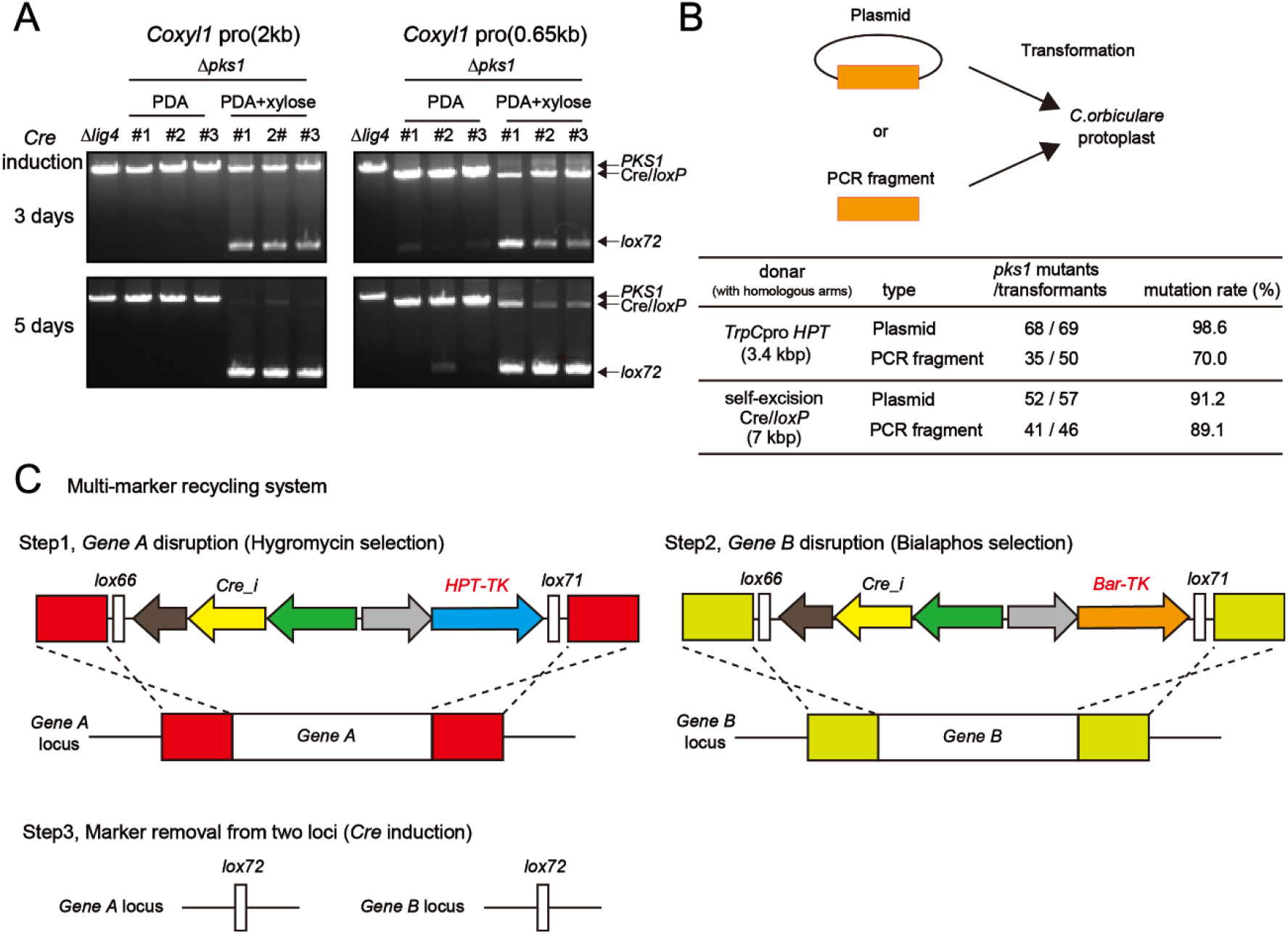
Improvement of self-excision Cre/*loxP* system. A, A short fragment of *Coxyl1* promoter also worked for Cre induction in the presence of xylose in *C. orbiculare*. B, Introduction of PCR fragments is sufficient to generate KO strains. PCR fragments were transformed with the CRISPR/Cas9 vector pChPtef026 containing *PKS1* gRNA1 into *C. orbiculare* protoplasts. C, A scheme of sequential gene disruption using multi-marker system with hygromycin-resistant gene and bialaphos-resistance gene. The double KO mutants were incubated on xylose media to remove selective marker cassettes from two loci.

For constructing targeting vectors, we fused a Cre/*loxP* fragment with homologous arms by PCR, and inserted into pCB1636 vector. We next tested whether this PCR fragment can directly be used as a DNA donar for gene targeting (Fig. 4B). We prepared two fragments, *TrpC* promoter *HPT* gene fragment (3.4 kbp) and Cre/*loxP* cassette (7 kbp), to investigate whether PCR fragment size affects gene-targeting efficiency; both fragments contain 1kbp 5’- and 3’-*PKS1* homology arms. We introduced these PCR fragments with the CRISPR/Cas9 vector pChPtef026 containing *PKS1* gRNA1 to *C. orbiculare* protoplasts. As a results, the direct introduction of a PCR fragment was sufficient to generate *pks1* mutants with high efficiency (Fig. 4B). In addition, high mutation rates were observed using both PCR fragments.

It takes more than 10 days for the marker recycling step including Cre induction on xylose media and negative selection for Cre/*loxP* cassette using FdU. To reduce the amount of time for multiple gene disruption, we also generated another self-excision Cre/*loxP* cassette using bialaphos resistance gene (*Bar*) (Fig. 4C). Multi-marker recycling system was reported to accelerate gene disruption in *Candida albicans* (*31*). Because multi-marker system was used for generating homozygous mutants in the diploid fungus *C. albicans* (*31*), we can establish double KO mutants in the haploid fungus *C. orbiculare* by using multi-marker system. After generating KO strains with hygromycin selection cassette, another gene can be sequentially targeted using bialaphos selection cassette. Cre can remove Cre/*loxP* cassettes from two loci in this double KO mutant.

### Sugar amounts in plants are altered during infection of *C. orbiculare*

Phytopathogens acquire host-derived sugars during infections (*32, 33*). We previously reported that *Arabidopsis* plants activate a sugar influx transporter to avoid pathogens’ sugar gain (*34*), revealing the importance of sugars in plant-pathogen interactions. In this study, to investigate what kinds of sugars *C. orbiculare* obtains from plants during infection, we attempted to identify sugar transporters involved in virulence of *C. orbiculare*. First of all, we measured sugar amounts in infected leaves. *C. orbiculare* is reported to infect not only *Cucumis sativus* but also *Nicotiana benthamiana* (*35*). Therefore, we monitored sugar amounts at 1, 3, 6 and 7 days post infection on true leaves of *N.benthamiana* and cotyledons of *C. sativus* (Fig. 5A and 5B). We found that quinate amounts increased in *N.benthamiana* leaves although it was reduced in cucumber cotyledons (Fig. 5C and 5D). On the other hand, cellobiose amounts were elevated in both plants during fungal infection (Fig. 5C and 5D).

**Fig. 5.**
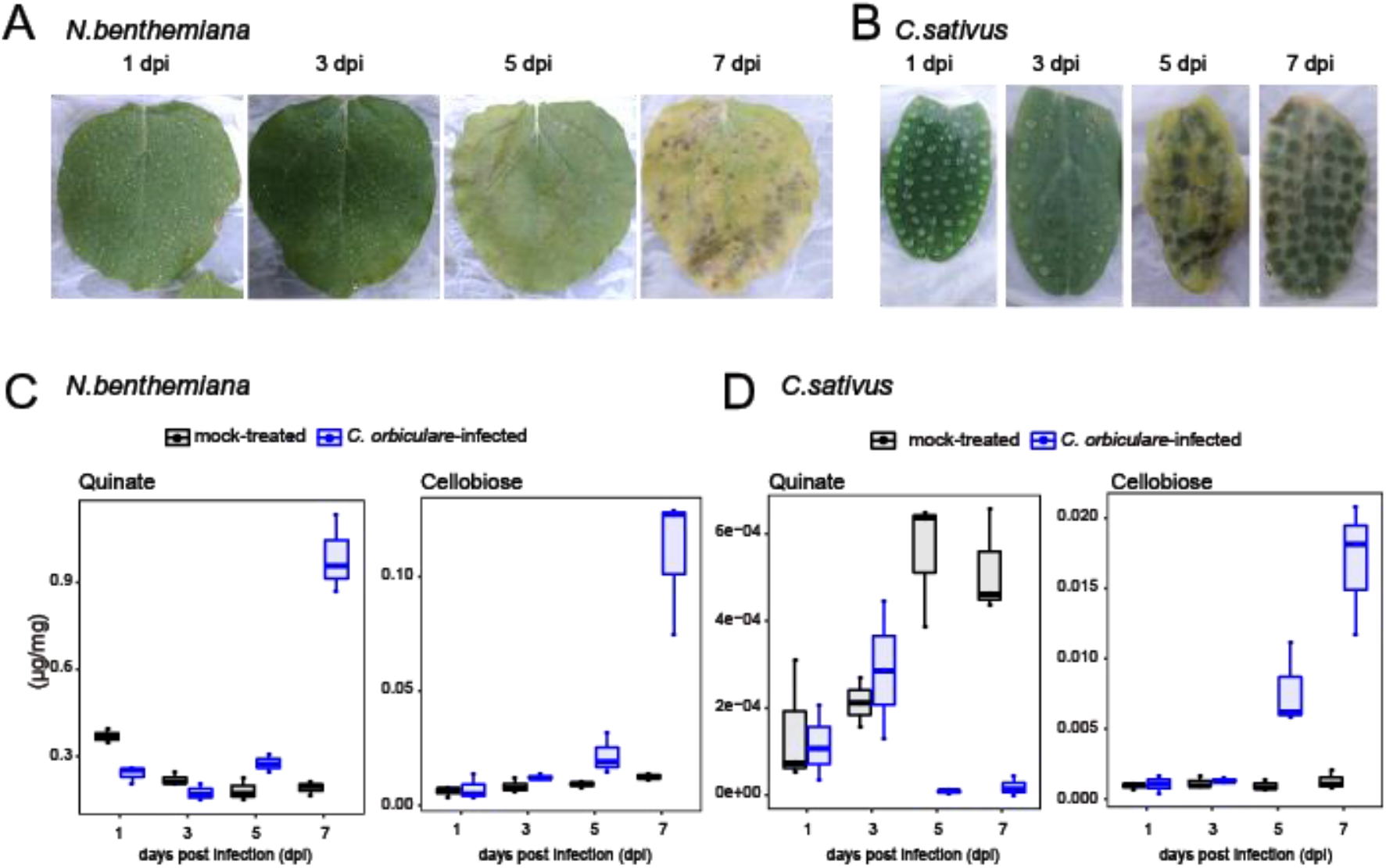
Sugar amounts in leaves are altered during the infection of *C. orbiculare*. A and B, The photograph of *N. benthamiana* leaves (A) and *C. sativus* cotyledons (B) during the infection of *C. orbiculare*. C and D, Quantification of quinate and cellobiose in *N. benthamiana* leaves (C) and *C. sativus* cotyledons (D) during the infection of *C. orbiculare*.

### Identification of quinate transporters in *C. orbiculare*

The expression of putative *C. orbiculare* quinate transporters was previously reported to be induced during the infection on *N. benthamiana* leaves (*35*). Therefore, we hypothesized that *C. orbiculare* acquires quinate during infection especially on *N.benthamiana*. To analyze this hypothesis, we investigated virulence of quinate transporter KO strains of *C. orbiculare*. The quinate transporter qa-y was identified in *N. crassa* (*36*). We found that five qa-y homologous genes in *C. orbiculare* (Fig. 6A). By using our self-excision Cre/*loxP* system, we established multiple knockout mutants which were deficient of five quinate transporters (Fig. 6B). Although mutant strains lacking *Cob_06838* and/or *Cob_05142* showed normal growth on minimal media containing quinate as a sole carbon source, the growth of the quadruple mutant defective of *Cob_06838, Cob_05142, Cob_05165* and *Cob_06690* and the quintuple mutant lacking all putative quinate transporters was impaired on quinate media. Importantly, the quadruple and the quintuple mutants normally grew on media containing glucose as a carbon source (Fig. 6C), confirming their functionality as quinate transporters in *C. orbiculare*. However, loss of these quinate transporters did not affect fungal virulence to *N. benthamiana* or *C. sativas* (Fig. 6E). We further found five other genes showing high homology with quinate transporters. These genes were homologs with the *N. crasse* galacturonic acid transporters GAT-1 (*37*) and the *Botrytis cinerea* hxt15 (*38*) (Fig. 6A). To further analyze the contribution of these transporter genes on virulence of *C. orbiculare*, we generated decuple mutants which are deficient of five quinate transporters and five galacturonic acid transporters (Fig. 6B). However, this strain showed normal growth on media containing galacturonic acid or pectin as a carbon source (Fig. 6D), indicating the presence of other galacturonic acid transporters in *C. orbiculare*. In addition, virulence of this decuple mutant was not altered on *N. benthamiana* or *C. sativus* (Fig. 6E).

**Fig. 6.**
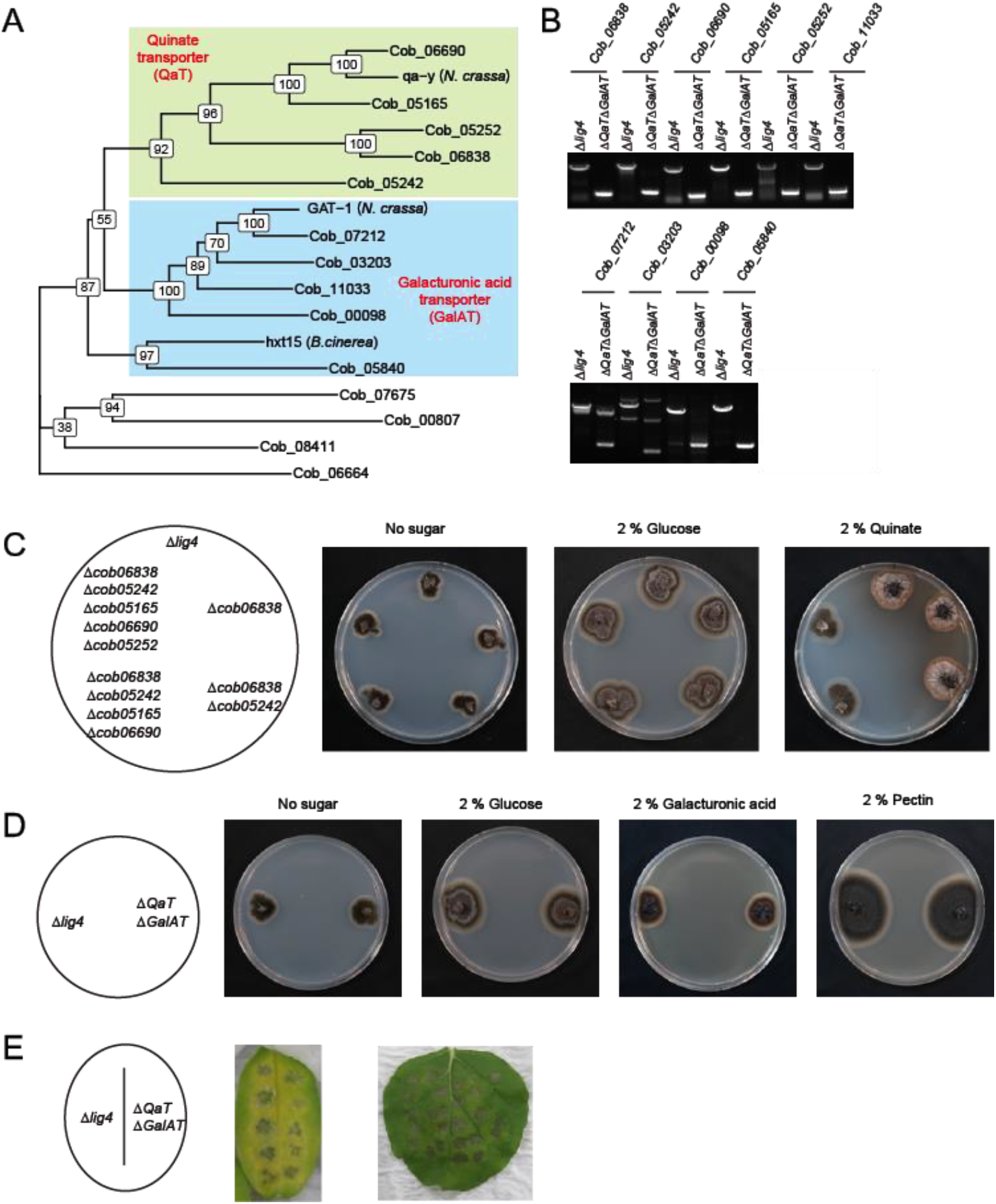
Identification of functional quinate transporters in *C. orbiculare*. A, A maximum-likelihood tree of quinate transporters (QaTs) and galacturonic acid transporters (GalATs) in *C. orbiculare* constructed by RaxML and ggtree is shown. Numbers on each node represent bootstrap values from 100 samplings. B, Genotyping of 5 *QaT* genes and 5 *GalAT* genes disrupted by self-excision Cre/*loxP* system. C, Loss of *QaT* genes restricted growth on quinate media, not glucose media. D, Loss of 5 *GalAT* genes did not affect growth on media containing galacturonic acid or pectin E, Loss of 5 *QaTs* and 5 *GalATs* did not affect virulence of *C. orbiculare*. The middle panel and the right panel shows cotyledon of *C. sativas* and true leaf of *N. benthamiana*, respectively. *lig4* mutant and *qat galat* mutant was inoculated on the left side and the right side, respectively, of infected leaves.

Because cellobiose amounts were elevated during infection on *N. benthamiana* and *C. sativus* (Fig. 5C and 5D), we also analyzed contribution of cellobiose transporters to virulence of *C. orbiculare*. We found that 26 genes which show higher homology with CDT-1 and CDT-2 which are previously characterized as cellobiose transporters in *N. crassa* (*39*) (Fig. 7A). Out of these 26 putative cellobiose transporters, 17 genes were knocked out using our self-excision Cre/*loxP* system in this study (Fig. 7B). This heptaducaple mutant showed normal virulence on *N. benthamiana* and *C. sativus* (Fig. 7C). In addition, this mutant strain was still able to grow on media containing cellobiose as a sole carbon source, suggesting redundant functions of remained 9 genes as cellobiose transporters (Fig. 7D).

**Fig. 7.**
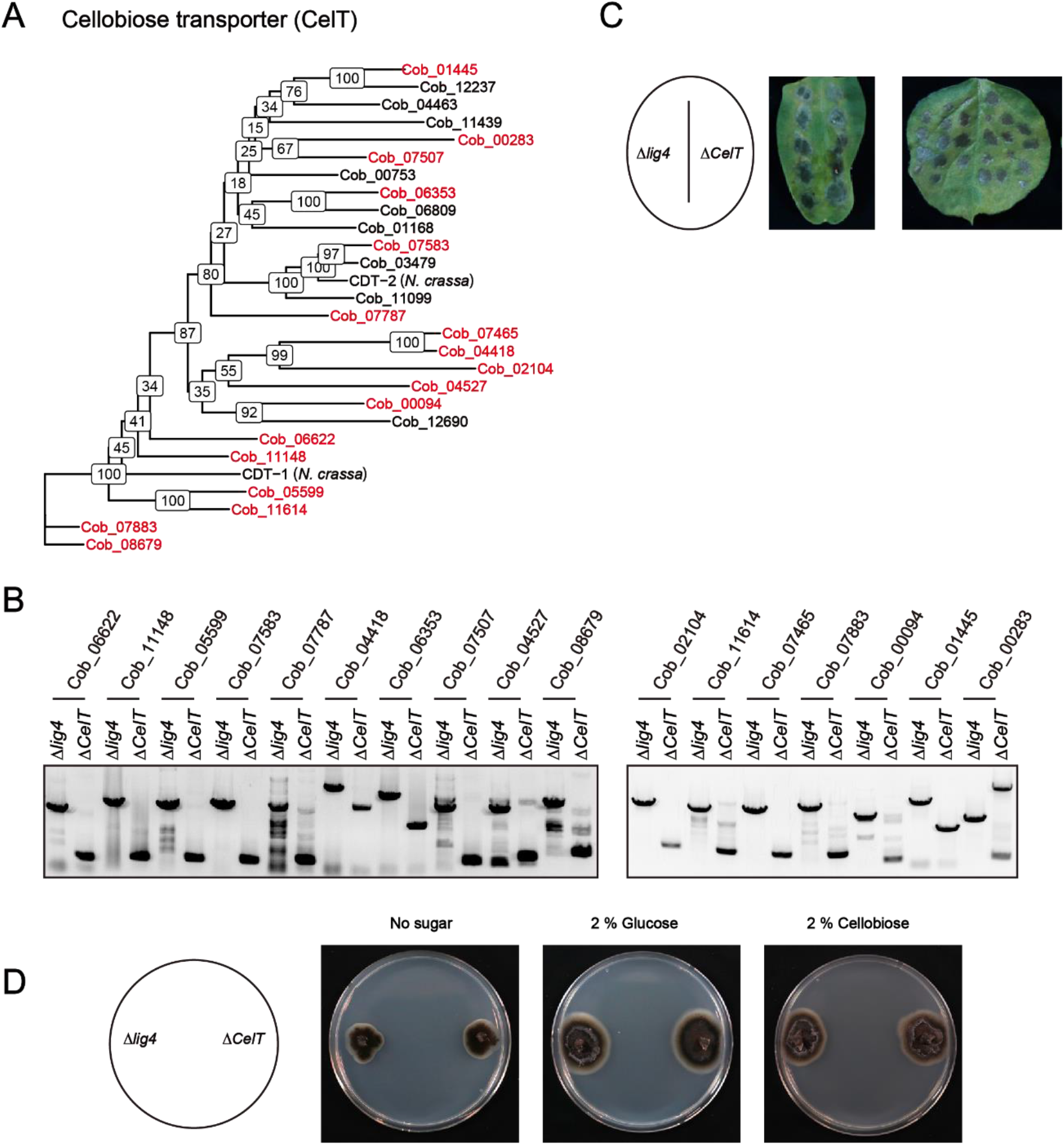
Disruption of cellobiose transporters in *C. orbiculare*. A, maximum-likelihood tree of cellobiose transporters (CelTs) in *C. orbiculare* constructed by RaxML and ggtree is shown. Numbers on each node represent bootstrap values from 100 samplings. Genes which are knocked out in this study were colored red. B, Genotyping of 17 *CelT* genes disrupted by self-excision Cre/*loxP* system. The marker cassette used for *Cob_00283* locus was not removed. C, Loss of 17 *CelT* genes did not affect virulence of *C. orbiculare*. The middle panel and the right panel shows cotyledon of *C. sativas* and true leaf of *N. benthamiana*, respectively. *lig4* mutant and *celt* mutant was inoculated on the left side and the right side, respectively, of infected leaves. D, Loss of 17 *CelT* genes did not affect growth on media containing cellobiose as a sole carbon source.

## Discussion

Although *C. orbiculare* has been employed as a model phytopathogenic fungus to analyze molecular mechanisms in plant-fungus interactions, sequential gene disruption was technically difficult. Because there are only a few desired selective markers for *C. orbuculare*, we applied Cre/*loxP*-based marker recycling system to this fungus in this study. Previously, the establishment of *URA3*-based marker recycling system was reported in *C. orbiculare* (*40*). The *URA3* gene encodes an orotidine-5-phosphate decarboxylase involved in uridine/uracil synthesis. *C. orbiculare* has two *URA3* genes, *URA3A* and *URA3B*, in the genome. The *ura3a/b* double mutants showed auxotrophy for uridine and insensitivity to 5-fluoroorotic acid (5-FOA). In background of the *ura3a/b* mutants, *URA3B* can be used as a selective marker for transformation under conditions without exogenous application of uridine because the introduction of *URA3B* provides prototrophy to *ura3a/b* mutants. Importantly, this *URA3B* selective marker cassette is removable when the transformants are incubated on PDA containing 5-FOA and uridine. However, because *ura3a/b* mutants lost virulence, *URA3B* gene needs to be re-introduced after *URA3B* removal step to analyze fungal virulence. As advantageous points of our Cre/*loxP* system from *URA3*-based system, our system is applicable to WT strains. In addition, we can prepare two positive selection markers, *HPT* and *Bar*. This multi-marker recycling system accelerates to generate multiple KO mutants by reducing the number of times of marker recycling steps. However, one *loxP* sequence is remained in the genome as a result of Cre-based recombination, possibly causing unfavorable recombination in the next excision rounds. To circumvent this risk, mutated *loxP* sequences, *lox66* and *lox77*(*25*), were used. After Cre-based recombination, inefficient *lox72* sequence is remained by using *lox66* and *lox77*. Here we disrupted 17 cellobiose transporter genes using this system, and did not observe any negative effects for fungal growth and virulence. These results suggested that unfavorable recombination caused by remained *lox72* sequences did not occur in our Cre/*loxP* system.

To obtain marker-free mutants, we also examined if CRISPR/Cas9-induced DSBs can directly introduce mutations on *PKS1* locus without donor DNAs because small insertions and/or deletions can be generated by the error-prone NHEJ pathway (*41*). However, we had never obtained *pks1* mutants without donor DNAs (data not shown). Likewise it was reported that this event rarely occur in *M. oryzae* (*42*). However, co-introduction of a telomere vector with CRISPR/Cas9 RNPs efficiently generated NHEJ-mediated mutations in *M. oryzae* (*43*). While such kinds of marker-free systems might be also developed for *C. orbiculare* near future, we thought that HR-based gene replacement with marker recycling systems is currently the most practical to build multiple mutations in *C. orbiculare*. Although disruption of NHEJ pathway massively promotes HR efficiency in *C. higginsianum* and other fungi (*4–6*), loss of *LIG4* only slightly enhanced it in *C. orbiculare* under our experimental conditions (Fig. 1D). However, gene-targeting efficiency using an inefficient gRNA was elevated in the absence of *LIG4* (Fig. 1D). Therefore, we employed the combination between CIRSPR/Cas9 system and *lig4* strain to generate KO strains in this study.

We previously described that plants activate a sugar influx transporter to inhibit pathogens’ sugar gain (*34*). Loss of sugar transporters causes enhanced susceptibility to pathogenic bacteria and fungi in *Arabidopsis* plants (*34, 44*), indicating that sugar uptake competition is important to shape plant-pathogen interactions. We found that quinate amounts were elevated during the infection of *C. orbiculare* in *N. benthamiana* leaves, but not in *C. sativa* cotyledons (Fig. 5C and 5D). These results suggested that *C. orbiculare* differently affects host sugar metabolism in a dependent manner on host plant species. During *M. orizae* infection, quinate amounts were also elevated in infected plants (*45*). *M. oryzae* might manipulate host quinate concentrations by modulating shikimate pathway to reduce defensive phenylpropanoid metabolism. In addition, quinate can be used as a carbon source by fungi such as *N. crassa* (*36*). Because we also showed that *C. orbiculare* can use quinate as a carbon source (Fig. 6C), we thought that *C. orbiculare* exploits elevated amounts of quinate as a carbon source during infection. However, loss of quinate transporters did not affect fungal virulence although it reduced fungal growth on media containing quinate as a sole carbon source (Fig. 6E). These results revealed that quinate uptake is dispensable for virulence of *C. orbiculare*.

We also found that cellobiose amounts increased during fungal infection in *N. benthemiana* and *C. sativus* (Fig. 5C and 5D). Cellobiose is generated by decaying cellulose, a major cell wall component. Leaves became necrotic during fungal infection (Fig. 5A and 5B), likely reflecting the amounts of cellobiose. Because many putative cellobiose transporters were found in the genome in *C. orbiculare*, this fungus may acquire cellobiose during infection. At least we showed that *C. orbiculare* can exploit cellobiose as a carbon source (Fig. 7D). Recently, the sugar transporter Hxk6 was identified to be involved in virulence of *C. higginsiaum* (*46*). Although ChHxk6 showed monosaccharide uptake activity in yeast, it was indeed a homolog of cellobiose transporters. Therefore, cellobiose uptake might be a key for virulence of *C. higginsianum* although cellobiose uptake activity of ChHxk6 was not analyzed yet. We here mutated Cob_07787 which is the closest homolog of ChHxk6 in *C. orbiculare*. However, the mutant strain did not show reduced virulence (Fig. 7C). Because the mutant was able to grow on media containing cellobiose as a sole carbon source (Fig. 7D), remained intact cellobiose transporters could redundantly work for cellobiose absorption. Therefore, further genetic study is required to analyze the importance of cellobiose uptake in fungal virulence. In addition, these results indicate the requirement of comprehensive gene-targeting approaches for loss-of-function study.

Phytopathogens have evolved various strategies to overcome plant immunity, including the use of effectors which suppress their host’s immune system. The genomes of many phytopathogenic fungi including *Colletotrichum* species have been reported, and these studies revealed the presence of multiple effector proteins (*35*). In most cases, effector proteins work redundantly to repress the same host proteins, indicating that multiple gene targeting methods are required to analyze their functions. In *U.maydis, FLP/FRT* system is applied to mutate multiple effector genes (*20*). We believe that CRISPR/Cas9 and Cre/*loxP* systems, we here developed, will become powerful tools for genetic studies to discover novel aspects in plant-fungus interactions.

## Materials and Methods

### Strains, culture conditions and infection assays

*C. orbiculare* strain 104-T (MAFF240422) was used as the WT strain in this study. *lig4* strain was previously established by disrupting *LIG4* gene in 104-T strain (*47*). Fungal strains were incubated on PDA medium (Nissui) or minimal medium (1.6 g/L yeast nitrogen base without amino acids (BD), 2 g/L asparagine (Wako), 1 g/L ammonium nitrate (Nacalai tesque), 15 g/L agar (Nacalai tesque)) containing 2 % indicated sugars at 24 °C under dark conditions. Selective agents were used at a final concentration of 100 μg/mL for hygromycin (Wako), 25 μg/mL for bialaphos (Wako), or 100 μM for 2’-deoxy-5-fluorouridine (FdU) (Tokyo Chemical Industry). For infection assays, true leaves from 3- or 4-week-old *N. bentehmiana* or cotyledons of 1-week-old *C. sativa* were used. Five μL of 5.0 x 10^5^ spores was dropped on detached leaves, and incubated in sealed dishes with wet paper for keeping humidity at 24 °C under 16 h light / 8 h dark cycle.

### Plasmid construction

The detailed information of plasmids used in this study are described in Fig. S2. For pChPtef026 vector, DNA fragments of the *C. higginsianum* U6 snRNA gene promoter with gRNA scaffold, the *A. melanogenum translation elongation factor 1-alpha* (*tef*) gene promoter, the fungal codon-optimized *Cas9* gene, and the *Agaricus bisporus heat shock protein 26 kDa* (*HSP26*) gene terminator, were synthesized by GenScript. Synthesized gene fragments were assembled and clone into the *Apa*I/*Eco*RV sites of pDONR221(Thermo Fisher Scientific) by using T4 DNA ligase (NEB) or In-Fusion HD-Cloning kit (Clontech). For inserting a gRNA fragment, annealed oligoDNA was ligated into *Bsa*I-digested pChPtef026 vector using T4 DNA ligase. For pChPtef026 Cre vector, fungal codon-optimized *Cre* gene, synthesized by FASMAC, was inserted to the *Nco*I/*Sac*I sites of pChPtef026 vector to replace *Cas9* with *Cre*. For pCB1636 *pks1* vector, 1 kbp upstream sequence from the start codon of *PKS1* and 1 kbp downstream sequence from stop codon of *PKS1* were assembled into the *Xho*I/*Sal*I sites and the *Cla*I/*Hin*dIII sites of pCB 1636 vector (*2*), respectively, by SLiCE reaction (*48*). For pCB1636 lox HPT-TK, DNA fragments of *lox66, TrpC* promoter, fungal codon-optimized *hygromycin phosphotransferase* (*HPT*) gene, fungal codon-optimized *thymidine kinase* (*TK*) gene and *lox71* were synthesized by FASMAC. These fragments were assembled to the *Sal*I/*Cla*I sites of pCB1636 vector by SLiCE reaction. For pCB636 *pks1* lox HPT-TK, 1 kbp upstream sequence from the start codon of *PKS1* and 1 kbp downstream sequence from stop codon of *PKS1* were assembled into the *Xho*I/*Sal*I sites and the *Cla*I/*Hin*dIII sites of pCB1636 lox HPT-TK vector, respectively, by SLiCE reaction. For pCB *pks1* clox HPT-TK, a *Coxyl1* promoter fragment was amplified from genome of *C. orbiculare. Cre, Coxyl1* promoter and *AbHSP26* terminator were assembled into the *Nhe*I/*Asc*I sites of pCB1636 *pks1* lox HPT-TK vector. For pCB1636 *pks1* clox Bar-TK, *HPT* gene of pCB *pks1* clox HPT-TK vector was replaced with bialaphos resistance (*Bar*) gene by PCR. For gene disruption of transporter genes, 1 or 0.5 kbp fragments of upstream from start codon and 1 or 0.5 kbp fragments of downstream from stop codon were amplified from genome of *C. orbiculare*. Self-excision Cre/*loxP* cassette was amplified from pCB1636 *pks1* clox HPT-TK vector or pCB1636 *pks1* clox Bar-TK vector. Three fragments (two homologous arms and one marker cassette) were fused by PCR (*49*). A fused DNA fragment was inserted into the *Xho*I/*Hin*dII sites of pCB1636 vector. All DNA fragments were amplified by KOD one (TOYOBO).

### Fungal transformation

PEG-mediated fungal transformation, which was previously described (*2*), was applied to introduce plasmids or PCR fragments to *C. orbiculare* protoplasts. Protoplasts were prepared by degrading cell wall via driselase (Sigma) and lysing enzyme (Sigma). Ten μg of plasmids or PCR fragments was used for transformation. Transformants were selected on PDA with 0.6 M glucose containing appropriate anti-biotics for 5-7 days. HR efficiency of *pks1* mutants was calculated at this time without PCR-based genotyping. Transformants were further transferred onto PDA medium with appropriate anti-biotics for several days. For genotyping PCR, mycelium was picked into 100 μL TE buffer and microwaved for 5 min twice. After centrifugation, supernatant was used as a template. For removing the selective marker cassette of self-excision Cre/*loxP* system, KO strains were transferred onto PDA medium containing 2 % xylose for 5 days. Conidia were harvested and streaked onto PDA media containing FdU. After a few days, single colonies were picked and transferred onto PDA medium containing FdU. For checking a removal of the selective marker cassette, genotyping of transformants were performed by the above-described method. In addition, transformants were transferred onto PDA medium containing appropriate selective agents to check whether they lost the resistance against the anti-biotics.

### Measurement of sugar content in infected leaves

*C. orbiculare*-infected leaves were frozen and ground in liquid nitrogen, and homogenized with 1 ml of extraction buffer (methanol:water:chloroform = 5:2:2) with 10 μg ribitol (Wako) as an internal control. Mixtures were incubated for 30 min at 37 °C. After centrifugation, 900 μl of the supernatant was transferred to a new tube, and 400 μl of water was added. After centrifugation, the upper phase was transferred to new tube. These samples were evaporated using a spin dryer at 50 °C, and subsequently freeze-dried. Samples were sonicated in 30 μl of methoxyamine (Sigma) (20 mg/ml dissolved in pyridine (Wako)), and incubated for 90 min at 30 °C. Subsequently, 30 μl of MSTFA + 1% TMCS (Thermo Fisher Scientific) was added and incubation was continued for 30 min at 37 °C. After centrifugation, the supernatants were subjected to gas chromatography-mass spectrometry analysis. Each sample (1 μl) was separated on a gas chromatograph (7820A; Agilent Technologies) combined with a mass spectrometric detector (5977B; Agilent Technologies). For quantitative determination of metabolites, peaks that originated from selected ion chromatograms (quinate 345, cellobiose 361, ribitol 319) were used.

## Acknowledgements

We thank Dr. Ryo Matsumoto (Tokushima University) for technical assistance, Prof. Yuriko Osakabe (Tokyo Institute of Technology) for discussion, and Prof. Katsuya Gomi (Tohoku University) for providing published materials. This work was supported by JST PRESTO (JPMJPR17Q9, K.Y.) and JSPS KAKENHI (20K21280, 21H02157, K.Y.; 21H05032, Y.T.).

## Author contributions

K.Y. and Y.T. conceived this study. K.Y. performed most experiments and data analyses. T. Y. and K.U. performed gene disruption analysis. K.O. provided plasmid vectors. K.Y. wrote the manuscript.

