## Supplemental figure for "The establishment of multiple knockout mutants of *Colletotrichum orbiculare* by CRISPR/Cas9 and Cre/*loxP* systems"

>*Coxyl1* promoter

ATCTCATCACCCCTCCGAGACGAGAGTCCGTACATGAGCACACTTTCGTTGGCTTGGGGTCCTGGCTTGTCGCAACCTT
CAGACCTTTGACGAACCCCAGCCAGAGTGAGTTTCGCAGCGCATGATTCGAGCGTCTGCCGGTCTCTTGACGTGAGAA
GGTGGCAGGGGCTCCCATGGAGCCGTCCTTTTAAGCTTGCTTGCTATTCGTGCTAGCAAGAATTTCGATGTCTGTTTTTG
GAGCACCATCCCTCTCACCAAAGTCTTCAGGTGATGGATGAAAGCGAGAACGTCTCAAACGCGCCGTGTTCTGGATCT
CGACGTTTTTTGTTTTCGCCAAGACGAACAGAAATATGGATTCCCCCAAGTCCAAATACTCCTATTTTCCGGACCATGAC
CAGTGTCCGGACTCAACTCTCACTCTGCCCATGGGCCATTGGTGCCACCACCAGTCTCCCAGGCAGGGACCTGATCAT
CACCGGAACATTTGTCATGTATGACTCGGGTAATAATGGACCTGTTTAAGGAGGATAAAAATATCGGAAACGACGCCTC
GATAAACGAATCACACATGCGAAACCTGCTGCACCTCCGGGCGCTGATTTTCCTCCTTGTTTCTGTTGCGTTCTTTGCTC
TCTGCTCGAACATTCTTCTAGCCATGATAACGGGCCCAAGCCTACCTCGCCGACTTGGCTCTTCATTAATCGTCCGTACT
ACGCTGAGCAGTGTCCTTCGTCCGGCACGTTTGCGACGAGGCCCCGACTTATCATCAGATGACGACATAGCCACACTC
ATTTTGGAATGCGGCCTCTTCTCAACACACACACACACACACACACACACACACTACTCTCCTATCGCTCGTGCTCGTA
ACAGCACGCCAGCTGATGACCACCCCTGCTCACTCCAACAGCTTCAGCCGTGCCTGTATCACCTCCGCGGCATGTTTTG
CCGTGAGATTGGACGGGATCGCTACCCAACCTCATAGCCGGTCGCAAGAATCCTCACAAGATCTGTGTTTTGAGAAGG
AATTTGCGACTTGGCGATCCTAGCATCAGGGCTGGCCGACGCTCGCGCGTTACCGAGTGTACGGTTTCCGGACATCTTC
TATTGTTGGAATTGCCAAGTCGCCTGCGGGCTCCGACGATCATATATTGCGACAGCAATTGGTTGCTTGCAAGAAAGGA
CTGACTCTGATCCCCGCGATCGGACAATCGGGTGAATCGTCGTCGCATGCAGTCCGAACTTGTAAGAAGAGGTTCCAC
CTCAGGACCTCTGCGTAGCGATAGTGAAGCGGGCGATCATGACTGCAGGGCTTGTTCGTCAGGTGGTTCAATGAGTGA
GAAGCAGGAAACTTGCCGGATTTACCCTTAGCAGGGTATTAAACAGCCTGAATGTGGATCATAATGCTTCACACAATGA
TCAGTTCTCCAGGAGCTACCACTAGGGACAAGTGCTGTTGCAGTCGCCATCTCGGCTGACGATGCCCTCATTCTTGCTT
GGTATCTATTCCTGAGTGTTCAACCTTGCGTACACCTTGCCCAGCACCGGTACCCTGGCAGGATAAAAGTAACTCGATC
GCATTTAACCGCTGCAAGAACGGGACGGGATGCAGGTCTTTACGAGCAGATGCCGTGACAGCCAACTACCGATGACCC
CAGTTCAAGTTGGTCTAGTGTACTCAAGGGGCCTATTTAGCCGGCCTCACATGTCACGAAGCCAAGTAATCACAACACT
CGATGCATCTCTCATCGCTGGCGCGCGGCATGCCATACCCGACTGGGTGTAACACTATGTTATGCAACAATGGTGAATTT
CGTTTTCCTCATCAGAGGTTTCTCGGAGCAATGGAGGTTTGGTCACGACTGACCTGCATTTTAGCCAGATACCTTGACA
GTGAGTAGCAGTAGAGAGCAATATTAAACCCGGAGATGTCGACAGTCCACTTGTACGAGCAACATCTTTGAATGCTATC
AACTTCATCATCGTCCCCTTACAATC

>Co*xyl2* promoter

GCCGGCGGCGCGATGGCGTGGAGGATCAACGCCGAGGACACGAGTGCCATGACGCAGCTCGCCATGAACTGGGGTCT
CAGCATCGGGTCGCTGATCCTCGCGCTGCCGACCGTGTTGACGGTGACGAAGAGCACGAGCGTCCTGGAGGAAGCCG
GCGCCGAGACGTCCAGTGAGAAAGCAAACGAGAAGATGGTGGAAGAATCATGACCTGTCGCACAGGTAGCAAGCGA
ATTCCTCGGTGATGCGGCTTGTATGCTACGTCCGATTTAATTACATGTACATTTTAGCAGGATCAACACCGAAAGAACTC
TGTACATACCTTACTTGGATGATTATTGCACCGTCACCCTTGAGGCTTGATGGCAGGAGACAATGATCGTCGAGTCGTG
ACACGAAGTCAAACCTCCTTTTGCTATTTCGTAGGAGCTGCCGCCAGATATAGTAGATCTAAACACCAGGAACTACTAG
GACTGATATAATAATCAAGGAATATGTTGATAGTTTAGTTTATCGTCATGGCGTGGTGAGCCACGAGCTGGCCCATACGT

TACCTATGGTTAAACCGACAGATTTTCAGATCGTTGCTTGGTGGCGAGTCCTCCTGCCATGCGAAAGAAACAAAGATAC

CCAGAAAGGCAGCTTGGAGCAGCTAGGATTTTTTTTTTCTAGCCCATGTCTGCCTAGCAGTCGTCACAATCAGCGGCTC

ACTGCTATGAAGCACCTGGTGGTAATTAGTTATGTGTCTACCTATCTCTGCTTGGCATTGATTCATTGACGCCTGGTGCTC

TGATCTGTGGCACGCTCGACCGCTTCATTCCGGCTCCTACCTTCTGACCAGCTGCCAGTGAGATTACAGGACACATTAC

TAATGACGAGTGTCACAATACCACTCTCAAAGTTTATTTCCCTCTTGTTGCGAAGGAGCCCACGTAGAAAGATGTGAAA

TCGTGTTGATGGAACAAAGGGATCTGGGGCGAAGAGGTTAAATCCCCCCTGAACCCACGCGAAGCCCAGACACGGGT

GGCTGTATGTGGCTTCGCAACACCTCCTACCAAGTAGTCTGATATGGAGGTATAATTCCACATTGTAGCCAAGGCCAAA

CAAACATGTACCTGGGCTATCTCTGGGCTTCGATCCCGGAACGTGGTCACTGTCCAGAAACCTTGACATTCCCGTCGTA

GCTGGCCACCACACCAGTGCTACGTCAAGTCTCGTGGTCCGCAGCCGGATATTCTCAAAGTGCATGCAGTGATCGGGA

GGCTCTATGGGAAATATCGAGACAGTAGCACAGCATGTTTTGCCGGCGGCTAAAAGGGGCGCTCCAGGTCCCCGGCGC

CGTGCTGTGACACAAAGTGTGGAGAAGCCGGTAGTCTTATGCGGCCTAGAACCTCCGAGTTTCTCTTCGTGGTTCGAG

TCTGCCGGTCATCCGATAGTTCAGATTCTAAGAAAGACGGGCTCGGTTTGGTGACGAGGGGCAGATGCGATGCTTTCA

GGATAGGCATCAGACCACGTGGGACTTTCGGTAGAAGCATGCGACTGTTTCATGTTTCGACGACGGAAAAGATTGAGC

TCTGGGTATTTGCGAGGCATTGATTAAGATCTACACAGTTAGTAGCCCATCTTATTGATACGAGAGAGCCATGAGGAACA

CCGCACTCGAATGCCCTTTTTCCATCTGGAACCCAGGTGCCAAGAATCATGTAACGGCTGAATGTAGAATACAACCCGA

TCAACAGACAGACCCCTTAGTGCAGCCACGGAAAGTCAGTTGCTGTGATGAAATGGTGTGCTGTTAATGAAGTTGCCA

TGAGTTGCCCCAATGGAGCAGTCTTCCGAGGTACCAGACCGAGTCTGACGATACTGTGAGACGTGGCGAACATCACGG

CCACAGGTCAATCGCATGCTGTCCGAGTATTGGGCGGTGTATAAAACGGCAGCCATATCGTCGTCCAACAATCGACAAT

CCTCCTCTCACCACCCACACTCCGACAACTGCAAAC

Figure S1, Promoter sequences of *Coxyl1* and *Coxyl2*.

2 kbp sequences from their stop codons of *Coxyl1* and *Coxyl2*. Putative XLR-1 binding sites, GGCTRR and GGNTAAAA, are colored yellow and green, respectively.

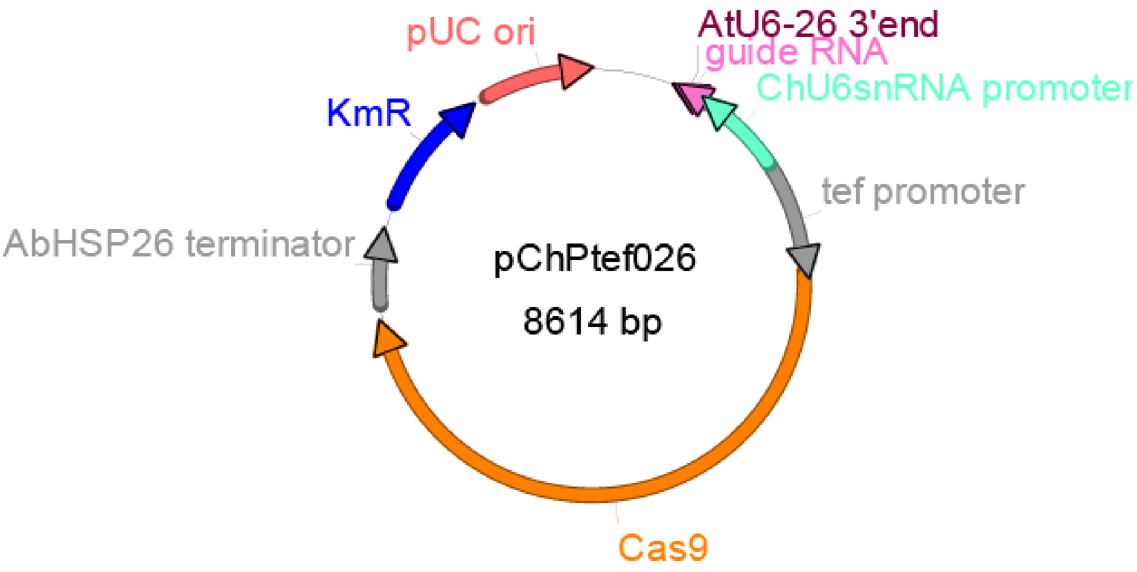


| Name | Location |
|---|---|
| AtU6-26 3'end | 618..625 |
| guide RNA | 626..702 |
| ChU6snRNA promoter | 727..1321 |
| tef promoter | 1330..2116 |
| Cas9 | 2119..6225 |
| AbHSP26 terminator | 6352..6831 |
| KmR | 7008..7817 |
| pUC ori | 7938..8611 |

>pChPtef026

CTTTCCTGCGTTATCCCCTGATTCTGTGGATAACCGTATTACCGCCTTTGAGTGAGCTGATACCGCTCGCCGCAGCCGAA

CGACCGAGCGCAGCGAGTCAGTGAGCGAGGAAGCGGAAGAGCGCCCAATACGCAAACCGCCTCTCCCCGCGCGTTG

GCCGATTCATTAATGCAGCTGGCACGACAGGTTTCCCGACTGGAAAGCGGGCAGTGAGCGCAACGCAATTAATACGCG

TACCGCTAGaCAGGAAGAGTTTGTAGAAACGCAAAAAGGCCATCCGTCAGGATGGCCTTCTGCTTAGTTTGATGCCTGG

CAGTTTATGGCGGGCGTCCTGCCCGCCACCCTCCGGGCCGTTGCTTCACAACGTTCAAATCCGCTCCCGGCGGATTTGT

CCTACTCAGGAGAGCGTTCACCGACAAACAACAGATAAAACGAAAGGCCCAGTCTTCCGACTGAGCCTTTCGTTTTAT

TTGATGCCTGGCAGTTCCCTACTCTCGCGTTAACgctaggATGGATGTTTTCCCAGTCACGACGTTGTAAAACGACGGCCA

GTCTTAAGCTCGGGCCCCGCTAGCAAGCTTTTTGAATTCCGCGGTGGCGCGGGATCCAACGGGAGCAAAAAAAAAGC

ACCGACTCGGTGCCACTTTTTCAAGTTGATAACGGACTAGCCTTATTTTAACTTGCTATTTCTAGCTCTAAAACGGAGAC

CGACAATTGAGGGTCTCCGAAAGCTGAGGAATGATTTGGTCACCCATGTATCCATCCATACCTTGGTTGAATGTCTGAC

GTGGCAGGTCTGTGCACCGTACTTGATGGGACTGCCCGAGGCGCCAGGTGCACCGCACCAGTCCTGAGTTCAAGCTCC

AGGCGCCCACCCCAACACAGAGGAGGATGACTGTCTGCCAGTGCTGTCAAAGTTTGTCGGTAGGGGATTCACCCCGTC

AAACGCACTGGGCCATCAGGAACTCGAAAACAAGGCAGGGGTTGATCTGATCTGTTCGGCCGTCGTTAAATAGTACGA

TCTTATTCCGAAAATCATCGTATTCCGTCAGCATCGCAACAAGTAGGCTTGGTCGGATCTCGGTGCACGGGATTTCCATC

CAGGTGCCGACTTGAGCAGTTTGTGTGGCGGTTCTGCAGATCAGGGGTTTCGAGTTGTGCGTAGGACGAACCTTTTGG

GTGACGGGAGAAATGGTGAGAAAAAGAATACAACAAAAAAGTTTTTTGGAAAATTTTGGCCGACGTTCTGAACCCCT

CCTTCCAGGCTTACCCCTCCTCCGAGAACCTCCTTTTCTGTTTCGTTTCGTTTTTGGCAATTCTTGGCGCGCCAAACGGT

GGTCAAAGGATGGTTCAGATACAAATTAGCAACAGGCCAGGCTAGACGCGCGACTATCCACTGCGGCAAATGGTGAGC

TGCAAGCAACGGTAAGATGTGACAGGACGAGCGGTGTGCCGGGAAAAAAATTGGAGGAGCGCAAAGCGGCGGCTGT

CCCTCAGTGGTGCCCAAACGTTATCGATAGTACACCAAGCATGGGCAGTGAGCGGCTATACAGAGGGAATAATAGGCAT

ATCGGCACGACTAGATTCGGTAGAAAGCATCGAAGAGCAATTCATTGAGCATATTATCACGTGGAATGCGATAGCTGTG

GCCAGGTTGAGACACCGCAAGTGAAAGATACACACATAGATTCTCGATTCGAGCGGTTTGCCTCCGCCACCGCAGTGC

ATAGCAAGCAAAGAAACGACAGTTGGCTCATCATCCGTTACATCATTTTTTCTACTGGCTCCGCTCGGTGGGCTCCCAA

CGAAGCAGCAAAAAAGTGAGAGAAAAAAACTAGCTTGGCGGGGCAACAGAAGCTAGACCCTTTGGCTCGCTTAGTCA

GTGCGCCCACTCACTCACACTCAAAAAGGCCACCCCTCCCGCACCCTCTTCTCATCACCGTCTTCATACCACGGTTCGT

CAAGCAATCGTATCTGGTAAGCTTTGACCTCCTCGAGCGGGCTCCACTTTGCTATTTCTTGGATCTGCTCTTTCTTTTCTC

TCTACCTCTTTTTCTAACCTCTCTTCAGAAAGTTCAACCGTACTTCACTCCATCTTCCATACATCACCGTCAAACCATGG

ACAAGAAGTACTCCATCGGCCTCGATATCGGCACCAACTCCGTCGGCTGGGCCGTCATCACCGACGAGTACAAGGTCC

CTTCCAAGAAGTTCAAGGTCCTCGGCAACACCGACCGCCACTCCATCAAGAAGAACCTCATCGGCGCCCTCCTCTTCG

ACTCCGGCGAAACCGCCGAGGCCACCCGCCTCAAGCGCACCGCCCGCCGCCGCTACACCCGCCGCAAGAACCGCATC

TGCTACCTCCAGGAAATCTTCTCCAACGAGATGGCCAAGGTCGACGACTCCTTCTTCCACCGCCTCGAGGAGTCCTTCC

TCGTCGAGGAGGACAAGAAGCACGAGCGCCACCCTATCTTCGGCAACATCGTCGACGAGGTCGCCTACCACGAGAAG

TACCCTACCATCTACCACCTCCGCAAGAAGCTCGTCGACTCCACCGACAAGGCCGACCTCCGCCTCATCTACCTCGCCC

TCGCCCACATGATCAAGTTCCGCGGCCACTTCCTCATCGAGGGCGACCTCAACCCTGACAACTCCGATGTCGACAAGC

TCTTCATCCAGCTCGTCCAGACCTACAACCAGCTCTTCGAGGAGAACCCTATCAACGCCTCCGGCGTCGACGCCAAGG

CCATCCTCTCCGCCCGCCTCTCCAAGTCCCGCCGCCTCGAGAACCTCATCGCCCAGCTCCCTGGCGAGAAGAAGAACG

GCCTCTTCGGCAACCTCATCGCCCTCTCCCTCGGCCTCACCCCTAACTTCAAGTCCAACTTCGACCTCGCCGAGGACGC

CAAGCTCCAGCTCTCCAAGGACACCTACGACGACGACCTCGACAACCTCCTCGCCCAGATCGGCGACCAGTACGCCG

ACCTCTTCCTCGCCGCCAAGAACCTCTCCGACGCCATCCTCCTCTCCGACATCCTCCGCGTCAACACCGAGATCACCAA

GGCCCCTCTCTCCGCCTCCATGATCAAGCGCTACGACGAGCACCACCAGGACCTCACCCTCCTCAAGGCCCTCGTCCG

CCAGCAGCTCCCTGAGAAGTACAAGGAAATCTTCTTCGACCAGTCCAAGAACGGCTACGCCGGCTACATCGACGGCG

GCGCCTCCCAGGAGGAGTTCTACAAGTTCATCAAGCCTATCCTCGAGAAGATGGACGGCACCGAGGAGCTTCTCGTCA

AGCTCAACCGCGAGGACCTCCTCCGCAAGCAGCGCACCTTCGACAACGGCTCCATCCCTCACCAGATCCACCTCGGCG

AGCTTCACGCCATCCTCCGCCGCCAGGAGGACTTCTACCCTTTCCTCAAGGACAACCGCGAGAAGATCGAGAAGATCC

TCACCTTCCGCATCCCTTACTACGTCGGCCCTCTCGCCCGCGGCAACTCCCGCTTCGCCTGGATGACCCGCAAGTCCGA

GGAAACCATCACCCCTTGGAACTTCGAGGAGGTCGTCGACAAGGGCGCCTCCGCCCAGTCCTTCATCGAGCGCATGAC

CAACTTCGACAAGAACCTCCCTAACGAGAAGGTCCTCCCTAAGCACTCCCTCCTCTACGAGTACTTCACCGTCTACAA

CGAGCTTACCAAGGTCAAGTACGTCACCGAGGGCATGCGCAAGCCTGCCTTCCTCTCCGGCGAGCAGAAGAAGGCCA

TCGTCGACCTCCTCTTCAAGACCAACCGCAAGGTCACCGTCAAGCAGCTCAAGGAGGACTACTTCAAGAAGATCGAG

TGCTTCGACTCCGTCGAGATCTCCGGCGTCGAGGACCGCTTCAACGCCTCCCTCGGCACCTACCACGACCTCCTCAAG
ATCATCAAGGACAAGGACTTCCTCGACAACGAGGAGAACGAGGACATCCTCGAGGACATCGTCCTCACCCTCACCCTC
TTCGAGGACCGCGAGATGATCGAGGAGCGCCTCAAGACCTACGCCCACCTCTTCGACGACAAGGTCATGAAGCAGCT
CAAGCGCCGCCGCTACACCGGCTGGGGCCGCCTCTCCCGCAAGCTCATCAACGGCATCCGCGACAAGCAGTCCGGCA
AGACCATCCTCGACTTCCTCAAGTCCGACGGCTTCGCCAACCGCAACTTCATGCAGCTCATCCACGACGACTCCCTCA
CCTTCAAGGAGGACATCCAGAAGGCCCAAGTCTCCGGCCAGGGCGACTCCCTCCACGAGCACATCGCCAACCTCGCC
GGCTCCCCTGCCATCAAGAAGGGCATCCTCCAGACCGTCAAGGTCGTCGACGAGCTTGTCAAGGTCATGGGCCGCCAC
AAGCCTGAGAACATCGTCATCGAGATGGCCCGCGAGAACCAGACCACCCAGAAGGGCCAGAAGAACTCCCGCGAGC
GCATGAAGCGCATCGAGGAGGGCATCAAGGAGCTTGGCTCCCAGATCCTCAAGGAGCACCCTGTCGAGAACACCCAG
CTCCAGAACGAGAAGCTCTACCTCTACTACCTCCAGAACGGCCGCGACATGTACGTCGACCAGGAGCTTGACATCAAC
CGCCTCTCCGACTACGACGTCGACCACATCGTCCCTCAGTCCTTCCTCAAGGACGACTCCATCGACAACAAGGTCCTC
ACCCGCTCCGACAAGAACCGCGGCAAGTCCGACAACGTCCCTTCCGAGGAGGTCGTCAAGAAGATGAAGAACTACTG
GCGCCAGCTCCTCAACGCCAAGCTCATCACCCAGCGCAAGTTCGACAACCTCACCAAGGCCGAGCGCGGCGGCCTCT
CCGAACTCGACAAGGCCGGCTTCATCAAGCGCCAGCTCGTCGAAACCCGCCAGATCACCAAGCACGTCGCCCAGATC
CTCGACTCCCGCATGAACACCAAGTACGACGAGAACGACAAGCTCATCCGCGAGGTCAAGGTCATCACCCTCAAGTCC
AAGCTCGTTTCCGACTTCCGCAAGGACTTCCAGTTCTACAAGGTCCGCGAGATCAACAACTACCACCACGCCCACGAC
GCCTACCTCAACGCCGTCGTCGGCACCGCCCTCATCAAGAAGTACCCTAAGCTCGAGTCCGAGTTCGTCTACGGCGAC
TACAAGGTCTACGATGTCCGCAAGATGATCGCCAAGTCCGAGCAGGAGATCGGCAAGGCCACCGCCAAGTACTTCTTC
TACTCCAACATCATGAACTTCTTCAAGACCGAGATCACCCTCGCCAACGGCGAGATCCGCAAGCGCCCTCTCATCGAA
ACCAACGGCGAGACTGGCGAGATCGTCTGGGACAAGGGCCGCGACTTCGCCACCGTCCGCAAGGTCCTCTCCATGCC
TCAGGTCAACATCGTCAAGAAGACCGAGGTCCAGACCGGCGGCTTCTCCAAGGAGTCCATCCTCCCTAAGCGCAACTC
CGACAAGCTCATCGCCCGCAAGAAGGACTGGGACCCTAAGAAGTACGGCGGCTTCGACTCCCCTACCGTCGCCTACTC
CGTCCTCGTCGTCGCCAAGGTCGAGAAGGGCAAGTCCAAGAAGCTCAAGTCCGTCAAGGAACTCCTCGGCATCACCA
TCATGGAGCGCTCCTCCTTCGAGAAGAACCCTATCGACTTCCTCGAGGCCAAGGGCTACAAGGAGGTCAAGAAGGAC
CTCATCATCAAGCTCCCTAAGTACTCCCTCTTCGAACTCGAGAACGGCCGCAAGCGAATGCTCGCCTCCGCCGGCGAA
CTCCAGAAGGGCAACGAACTCGCCCTCCCTTCCAAGTACGTCAACTTCCTCTACCTCGCCTCCCACTACGAGAAGCTC
AAGGGCTCCCCTGAGGACAACGAGCAGAAGCAGCTCTTCGTCGAGCAGCACAAGCACTACCTCGACGAGATCATCGA
GCAAATCTCCGAGTTCTCCAAGCGCGTCATCCTCGCCGACGCCAACCTCGACAAGGTCCTCTCCGCCTACAACAAGCA
CCGCGACAAGCCTATCCGCGAGCAGGCCGAGAACATCATCCACCTCTTCACCCTCACCAACCTCGGCGCCCCTGCCGC
CTTCAAGTACTTCGACACCACCATCGACCGCAAGCGCTACACCTCCACCAAGGAGGTCCTCGACGCCACCCTCATCCA
CCAGTCCATCACCGGCCTCTACGAAACCCGCATCGACCTCTCCCAGCTCGGCGGCGACGGATCTGCTGACGGTCCTTC
ACTGGGTTCAGGGTCACCCAAGAAGAAACGCAAAGTCGAAGATCCAAAGAAGAAAAGGAAGGTTGAAGACCCCAAG
AAAAAGAGGAAGGTGGATGAGTTCTAGGAGCTCTTAGTAGCTCCTCGGTTTTGGACTTTTATTTCGATTGTTTTAAATG
GCCTTGTTTTATGCATTGCGAGTTTAGCTTTCTTGGTCGTCATTATATGATTATTTTTTGATAGAAACACTTTCACCTCCCC
TCTTGCACATGTTGTATCCATTATGATGTATTTTATCGGACTCGACTCCCTACTTCAGAGTATTCTTTTATTTTCTCTAGCAT
CTTGTAATCGTAAATAATACCTTTCTTAATCTGCATTTACCATTACCAAATTTACCTCCTAAATGGTCCTTCCGCACTCAA

AGTGAAATGGCTGTACGACTCAAGTTCCGCAGTACAGGGCTGTTCCGGATAGACCACACTGTACCGCACGAAGCACAT

CTGAAATTTACCTTCGCGTACTAGACACCCCAAATAGTACGCTCCACGGCTTAAGCCTTAACATGCCTTGTGATTCTCTT

CTAAGTTATAAATAACGCAATGCTGTTCAATACGTACCTTAATTAAGTTTATCCCCTATAGTGAGTCGTATTACATGGTCAT

AGCTGTTTCCTGGCAGCTCTGGCCCGTGTCTCAAAATCTCTGATGTTACATTGCACAAGATAAAAATATATCATCATGAA

CAATAAAACTGTCTGCTTACATAAACAGTAATACAAGGGGTGTTATGAGCCATATTCAACGGGAAACGTCGAGGCCGCG

ATTAAATTCCAACATGGATGCTGATTTATATGGGTATAAATGGGCTCGCGATAATGTCGGGCAATCAGGTGCGACAATCT

ATCGCTTGTATGGGAAGCCCGATGCGCCAGAGTTGTTTCTGAAACATGGCAAAGGTAGCGTTGCCAATGATGTTACAGA

TGAGATGGTCAGACTAAACTGGCTGACGGAATTTATGCCTCTTCCGACCATCAAGCATTTTATCCGTACTCCTGATGATG

CATGGTTACTCACCACTGCGATCCCCGGAAAAACAGCATTCCAGGTATTAGAAGAATATCCTGATTCAGGTGAAAATAT

TGTTGATGCGCTGGCAGTGTTCCTGCGCCGGTTGCATTCGATTCCTGTTTGTAATTGTCCTTTTAACAGCGATCGCGTATT

TCGTCTCGCTCAGGCGCAATCACGAATGAATAACGGTTTGGTTGATGCGAGTGATTTTGATGACGAGCGTAATGGCTGG

CCTGTTGAACAAGTCTGGAAAGAAATGCATAAACTTTTGCCATTCTCACCGGATTCAGTCGTCACTCATGGTGATTTCT

CACTTGATAACCTTATTTTTGACGAGGGGAAATTAATAGGTTGTATTGATGTTGGACGAGTCGGAATCGCAGACCGATAC

CAGGATCTTGCCATCCTATGGAACTGCCTCGGTGAGTTTTCTCCTTCATTACAGAAACGGCTTTTTCAAAAATATGGTAT

TGATAATCCTGATATGAATAAATTGCAGTTTCATTTGATGCTCGATGAGTTTTTCTAATCAGAATTGGTTAATTGGTTGTA

ACACTGGCAGAGCATTACGCTGACTTGACGGGACGGCGCAAGCTCATGACCAAAATCCCTTAACGTGAGTTACGCGTC

GTTCCACTGAGCGTCAGACCCCGTAGAAAAGATCAAAGGATCTTCTTGAGATCCTTTTTTTCTGCGCGTAATCTGCTGC

TTGCAAACAAAAAAACCACCGCTACCAGCGGTGGTTTGTTTGCCGGATCAAGAGCTACCAACTCTTTTTCCGAAGGTA

ACTGGCTTCAGCAGAGCGCAGATACCAAATACTGTTCTTCTAGTGTAGCCGTAGTTAGGCCACCACTTCAAGAACTCTG

TAGCACCGCCTACATACCTCGCTCTGCTAATCCTGTTACCAGTGGCTGCTGCCAGTGGCGATAAGTCGTGTCTTACCGG

GTTGGACTCAAGACGATAGTTACCGGATAAGGCGCAGCGGTCGGGCTGAACGGGGGGTTCGTGCACACAGCCCAGCT

TGGAGCGAACGACCTACACCGAACTGAGATACCTACAGCGTGAGCTATGAGAAAGCGCCACGCTTCCCGAAGGGAGA

AAGGCGGACAGGTATCCGGTAAGCGGCAGGGTCGGAACAGGAGAGCGCACGAGGGAGCTTCCAGGGGGAAACGCCT

GGTATCTTTATAGTCCTGTCGGGTTTCGCCACCTCTGACTTGAGCGTCGATTTTTGTGATGCTCGTCAGGGGGGCGGAG

CCTATGGAAAAACGCCAGCAACGCGGCCTTTTTACGGTTCCTGGCCTTTTGCTGGCCTTTTGCTCACATGTT

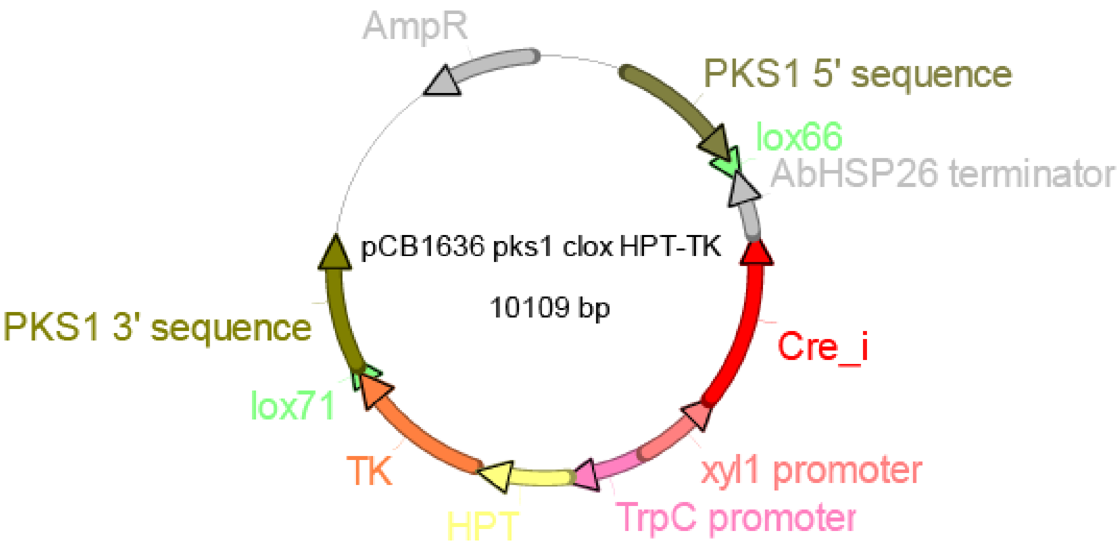


| Name | Location |
|---|---|
| PKS1 5' sequence | 675..1675 |
| lox66 | 1715..1748 |
| AbHSP26 terminator | 1760..2239 |
| Cre_i | 2240..3580 |
| xyl1 promoter | 3581..4232 |
| TrpC promoter | 4233..4917 |
| HPT | 4918..5586 |
| TK | 5623..6753 |
| lox71 | 6763..6796 |
| PKS1 3' sequence | 6837..7836 |
| AmpR | 9121..9981 |

>pCB1636 pks1 clox HPT-TK

CTGACGCGCCCTGTAGCGGCGCATTAAGCGCGGCGGGTGTGGTGGTTACGCGCAGCGTGACCGCTACACTTGCCAGCG

CCCTAGCGCCCGCTCCTTTCGCTTTCTTCCCTTCCTTTCTCGCCACGTTCGCCGGCTTTCCCCGTCAAGCTCTAAATCGG

GGGCTCCCTTTAGGGTTCCGATTTAGTGCTTTACGGCACCTCGACCCCAAAAAACTTGATTAGGGTGATGGTTCACGTA

GTGGGCCATCGCCCTGATAGACGGTTTTTCGCCCTTTGACGTTGGAGTCCACGTTCTTTAATAGTGGACTCTTGTTCCAA

ACTGGAACAACACTCAACCCTATCTCGGTCTATTCTTTTGATTTATAAGGGATTTTGCCGATTTCGGCCTATTGGTTAAAA

AATGAGCTGATTTAACAAAAATTTAACGCGAATTTTAACAAAATATTAACGCTTACAATTTGCCATTCGCCATTCAGGCT

GCGCAACTGTTGGGAAGGGCGATCGGTGCGGGCCTCTTCGCTATTACGCCAGCTGGCGAAAGGGGGATGTGCTGCAA

GGCGATTAAGTTGGGTAACGCCAGGGTTTTCCCAGTCACGACGTTGTAAAACGACGGCCAGTGAGCGCGCGTAATACG

ACTCACTATAGGGCGAATTGGGTACCGGGCCCCCCCTCGAGTGCCTACAAACCTGCCGTATACGGGCCATATTCGCGGG

CAGAAGGGCTCCGCCGACTCGATAATCCTGTGTGACCCTCGCGGCATCAGCGTCACGTCAACATCTAGCCTCGTTGGA

CTGGGGATGCTATGATATCAATGCATGTAGTGGACAAGCTCACGCAATGCTTACCTGCAGTTACAGTCCCTGCTGCGCG

GTGTCGAGAAGCCCGAAGCCTGGGTACGTGATACGGTACCTCGCATGCGCACCCTCGCTTCTGCTGGCTGGCGTTTTGT

TATTCTGGGCGTCTGGGATCAGAGCCAGGTAAGAAAGATGGGAAGAATGAGTGAGCACGCCTTGTGCTTTGCTCTGCC

TTGCTGCTGCGCTAGACATGCTTTGCCGTGCCGTGCTGCTGCCCTGCGCTGCGGCATGCTGCGATCGTTCCTCTTGTCTG

TCCGCCTGTCCGTCCTCCACCCCGGGCCACGATAGAGGAGCTGGCGCAAGGAAGCCCGTGTGGCGGCCTGGATCGCCT

GCCCTGAACCAATGGGTCTCGACCGATGCGGCAAGACAGCGAAAATCATCCATCAGCTTCGGTCACCGAGGAAGCTCA

GGGCCGGACTCACGCTATCCACCCGGGCAAGAAAAGGATGGGCCAAGTCAGTGGGAAAGGAAGGGGTGCAAGGGGG

GCGTGCGGATACATGGGACGTCTTCACCACACCCCCTTCTCCTTCTCTCGTTCTCACCTCACTCCCTCGACCCAGACCA

CTTGGGTTGTGTGTTCCATGTTGCAATTCCGCTATCCATCAGCCATGACTTTGACAGATGGGTTTGATTATTTTGGGGAG

ACGCCAGCTCGTGGTGGTGTTGACCTTGTCCATGGTGTGTACACCCCGTCTGTCCTTTGGCCCATCTCTTTCATATAAGT

TCCAGACTCGCTCCCAGTATCCCAATCTCAGTTTTTGTTAGGACACCGGGAAACCTCTCAAACACCTAGTAGGAAATTC

ATGTCGACAACCTGACAAGGTCGACGTTAACTGATATTGAAGGAGCATTTTTTGGGCTTACCGTTCGTATAGCATACATT

ATACGAAGTTATTGGGGCGCGCCTTGAACAGCATTGCGTTATTTATAACTTAGAAGAGAATCACAAGGCATGTTAAGGC

TTAAGCCGTGGAGCGTACTATTTGGGGTGTCTAGTACGCGAAGGTAAATTTCAGATGTGCTTCGTGCGGTACAGTGTGG

TCTATCCGGAACAGCCCTGTACTGCGGAACTTGAGTCGTACAGCCATTTCACTTTGAGTGCGGAAGGACCATTTAGGAG

GTAAATTTGGTAATGGTAAATGCAGATTAAGAAAGGTATTATTTACGATTACAAGATGCTAGAGAAAATAAAAGAATACT

CTGAAGTAGGGAGTCGAGTCCGATAAAATACATCATAATGGATACAACATGTGCAAGAGGGGAGGTGAAAGTGTTTCT

ATCAAAAAATAATCATATAATGACGACCAAGAAAGCTAAACTCGCAATGCATAAAACAAGGCCATTTAAAACAATCGAA

ATAAAAGTCCAAAACCGAGGAGCTACTAAGAGCTCTTAATCGCCGTCTTCGAGGAGACGGACCATAGCGCCGGTCTCG

CTGTCGAGGTTGCGGATGTAGTTCATGACGATGTTGACGTTGGTCCAGCCTCCGGCTTGCATGATCTCAGGAATGGAGA

CGCCGGCGCGGGCCATGTCGCGGGCGGCGCCCACGCGGGCGGAGTGGCCGGACCAGGCGAGGTAGCGCTGGCCAGA

ATCGTCCTTGGCACCGTAGATCAGGCGATGGGTGGCTTCGAAGATACCCTCCAGGGCGCGGGTCGAGAGTTGGGACGT

GGCGCTAGGGGCAGCGACGCCGTTCTTGCGCACGCGACAGAAGAGGTAGTTGTTAGGGTCGTCGGCAACGCCCGAAA

CGCTAATCCAGCGTTCGACGAGCTTAGTGACACCGAGGCTGAGCGCCTTTTCCACACCAGCTGTGCTCACCAAAGTTT

TGGTTCTTCCGATGTGAATGAGCATGCCTGAAAAAGATCAGTACGATGTTTGATAGAATGAAATAGGAAGTCGTGGTGA

AGCCCGGGTTAAGCCTGGGAAGACGCGTCTAGATGCGTGATGCATGCCAACGGGCAAGTGCCCATTACACGGCGGGGT

ACGCCACGATGCGAGATGCGAGACGCGCGCCAAAACCGACAGCGACGACGAGGTATAATAAAAATGGTACTTACGTCC

GCCATCGGTTCGGCTGATGTCCTTCACGCGAATGCGCGCGATTTCGGCGATGCGGAGCAAGGTGTTGTACGCGATACCC

AGAAAAGCAAGATTTCGGATGTCCTGGCAGCGGTCCGAGTTCTCCATAAGAGATCGGACCTGGTCGAAATCCGTTCTC

TCGAAAGCCAGAGCTTGCTTAGCTCGTTCTCCCGCATCGACGTTTTCTTTTCGGATTCGTCTCATAACCAGCGAGACCG

CATTCGAGTCGCTGGGTCGGGGAAGACCCGAACGTCTGTGAAGCATGTTCAACTGTCCCAAGTGCTGCTGAATGGTCT

TGACAGCCAGGCCTCTAGCCTGCAAATACAGCAGATAGTCACGGACGTCCTCTGGCTCGGCAGGAAACCACTTTCGGT

TGTTCAACTTGCACCATGCTGCCCAGGAACGGCAGACGCTCAGGAGCATCTTCCAAGTATGCTCGGAAAAGGCTTGGC

GATCTCGGAACATGTCCATCAAGTTCTTTCGGACTTCATCGGAGGTTGCATCAACCGGCAGGGCCGGGAGATTCTGGTG

GACTGTCAACAGGTTGCTCTCAAGCAGCTCCTTCAGGTGCGCCGCCTTTCGATAACGGCTACGTCTGGAGGACTCTCTA

TTCGACTCCTTTCTCTTACGAACGCGCTCTTCCGTTGCCATGGGATTGTAAGGGGACGATGATGAAGTTGATAGCATTCA
AAGATGTTGCTCGTACAAGTGGACTGTCGACATCTCCGGGTTTAATATTGCTCTCTACTGCTACTCACTGTCAAGGTATC
TGGCTAAAATGCAGGTCAGTCGTGACCAAACCTCCATTGCTCCGAGAAACCTCTGATGAGGAAAACGAAATTCACCAT
TGTTGCATAACATAGTGTTACACCCAGTCGGGTATGGCATGCCGCGCGCCAGCGATGAGAGATGCATCGAGTGTTGTGA
TTACTTGGCTTCGTGACATGTGAGGCCGGCTAAATAGGCCCCTTGAGTACACTAGACCAACTTGAACTGGGGTCATCGG
TAGTTGGCTGTCACGGCATCTGCTCGTAAAGACCTGCATCCCGTCCCGTTCTTGCAGCGGTTAAATGCGATCGAGTTAC
TTTTATCCTGCCAGGGTACCGGTGCTGGGCAAGGTGTACGCAAGGTTGAACACTCAGGAATAGATACCAAGCAAGAAT
GAGGGCATCGTCAGCCGAGATGGCGACTGCAACAGCACTTGTCCCTAGTGGTAGCTCCTGGAGAACTGATCATTGTGT
GAAGCATTATGATCCACATTCAGGCTGTTTAATACCCTGCTAAGGGTAAATCCGGCAAGTTTGCTAGCGAGCTAGTGGA
GGTCAACAATGAATGCCTATTTTGGTTTAGTCGTCCAGGCGGTGAGCACAAAATTTGTGTCGTTTGACAAGATGGTTCA
TTTAGGCAACTGGTCAGATCAGCCCCACTTGTAGCAGTAGCGGCGGCGCTCGAAGTGTGACTCTTATTAGCAGACAGG
AACGAGGACATTATTATCATCTGCTGCTTGGTGCACGATAACTTGGTGCGTTTGTCAAGCAAGGTAAGTGGACGACCCG
GTCATACCTTCTTAAGTTCGCCCTTCCTCCCTTTATTTCAGATTCAATCTGACTTACCTATTCTACCCAAGCATCCAAATG
AAAAAGCCTGAACTCACCGCGACGTCTGTCGAGAAGTTTCTGATCGAAAAGTTCGACAGCGTCTCCGACCTGATGCA
GCTCTCGGAGGGCGAAGAATCTCGTGCTTTCAGCTTCGATGTAGGAGGGCGTGGATATGTCCTGCGGGTAAATAGCTGC
GCCGATGGTTTCTACAAAGATCGTTATGTTTATCGGCACTTTGCATCGGCCGCGCTCCCGATTCCGGAAGTGCTTGACAT
TGGGGAGTTCAGCGAGAGCCTGACCTATTGCATCTCCCGCCGTGCACAGGGTGTCACGTTGCAAGACCTGCCTGAAAC
CGAACTGCCCGCTGTTCTCCAGCCGGTCGCGGAGGCCATGGACGCAATCGCCGCCGCCGACCTCTCCCAAACTTCCGG
ATTTGGCCCTTTCGGTCCCCAGGGCATTGGCCAGTACACTACCTGGCGAGACTTCATCTGTGCCATCGCAGATCCCCAC
GTTTATCATTGGCAGACCGTGATGGATGACACCGTGTCGGCCAGCGTCGCACAGGCCTTGGATGAGCTTATGCTGTGGG
CCGAAGATTGCCCTGAGGTTAGACATCTGGTTCATGCGGATTTCGGCAGCAACAATGTCCTTACTGATAACGGTCGCAT
CACGGCCGTCATTGACTGGTCCGAGGCGATGTTCGGCGATTCTCAGTATGAAGTCGCAAATATCCTCTTTTGGCGTCCTT
GGCTGGCTTGTATGGAACAGCAGACGCGTTACTTCGAGCGCAGACACCCTGAGCTGGCAGGATCCCCAAGACTGCGA
GCCTACATGCTCCGAATTGGACTCGACCAGCTCTATCAGTCCCTCGTGGATGGCAACTTTGACGATGCGGCGTGGGCTC
AAGGCCGTTGCGACGCCATCGTGCGATCCGGAGCAGGCACTGTCGGCCGCACACAGATCGCTCGCAGATCTGCCGCTG
TCTGGACTGACGGATGCGTGGAGGTTCTCGCGGATTCCGGTAACCGCAGACCCTCCACTCGCCCACGCGCCAAAGAAA
TCGATGGAGCTGGAGCAGGAGCTGGAGCCGGTGCGATGGCTTCCTACCCCTGCCATCAGCACGCCTCCGCCTTTGACC
AGGCGGCTAGATCCCGCGGACATAGCAACCGCAGAACAGCCCTCCGACCGCGCCGTCAACAGGAAGCGACGGAAGTT
CGATTGGAGCAGAAGATGCCCACCCTCCTTAGAGTCTATATCGACGGCCCTCACGGAATGGGCAAGACGACCACCACC
CAGCTCCTCGTCGCACTGGGTAGCCGTGATGACATCGTCTACGTTCCTGAGCCAATGACGTATTGGCAAGTGCTCGGCG
CATCCGAAACCATTGCCAACATTTACACCACCCAACACCGCCTGGACCAGGGTGAAATTTCCGCGGGAGATGCTGCCG
TCGTCATGACTTCGGCCCAAATTACTATGGGTATGCCGTACGCCGTCACGGACGCTGTCCTTGCACCCCACGTGGGCGG
AGAGGCCGGTTCGTCTCATGCCCCTCCTCCTGCCCTCACCCTTATCTTCGATCGCCACCCTATTGCCGCGTTGCTCTGTT
ACCCTGCTGCCCGCTACCTCATGGGTTCTATGACACCGCAGGCTGTCCTTGCTTTCGTCGCTCTCATCCCGCCAACACT
GCCCGGCACTAATATCGTCCTTGGCGCTCTCCCAGAAGACCGACACATTGATAGACTCGCCAAGCGCCAGAGACCTGG
AGAGAGATTGGACTTGGCTATGCTCGCCGCCATCCGTCGTGTCTACGGCCTCCTCGCTAACACAGTCAGATATCTGCAG

GGAGGCGGATCCTGGTGGGAGGACTGGGGTCAGCTTTCGGGTACTGCTGTGCCTCCCCAGGGCGCCGAGCCGCAGAG

CAACGCCGGTCCCAGACCCCATATCGGTGACACGCTCTTTACCCTGTTCCGCGCGCCTGAGCTGCTGGCCCCGAACGG

AGATTTGTACAACGTCTTCGCCTGGGCCTTGGACGTGCTCGCTAAGCGACTCCGCCCAATGCACGTGTTCATCTTGGAC

TACGACCAATCTCCTGCCGGTTGCCGCGACGCCCTCCTGCAATTGACCTCTGGCATGGTTCAGACCCACGTCACTACAC

CGGGAAGCATCCCTACGATCTGCGATCTTGCCCGCACATTCGCACGAGAGATGGGCGAAGCGAATTAAAGTCATATGAT

AACTTCGTATAGCATACATTATACGAACGGTAAGATGCCGACCGGGAACCAGTTAACGTCGACGGTATCGATGCCAAGC

AACTTGGAACTTTCCTTCGTGAAGCGGTTTGCGAGTAAGGGATTCACATAGCGAGGTTGCATACAAGCACGCGCATAG

CAGGTATAGAAGGGAAACAGGACATTACATGATTGCGAGCATGGATATGAGACTGCCTCTTTTTTTCCATCATTGTTCAT

TTTTCTTGTCATCTTTTCATCTTTCTTTTTTACCACCCTGTCTTAACATTGATATCCGCTCCCTGTACAAGCGCGAGATCAT

GCACACTCTTACGAATCATTCCTTACTCAACGTCGGCCGGAAAGGGAGGTAGAGGTGGAATTATCTGGGGAAACAGCA

TGGTCAACACCTCAGCCTTCGACCTTCTTGGGCACGGGAAGGACGGATGGCGCGGGTGTTGGCTTCGAAAGTAATTGG

GCTCCATCATTTTCTTTTGTCACGAATTAGCAGATTCCTCATGTCAGGCGGTGTTTACCAGGTTTTGTCGTCTTTTTCTGC

GGTCCATAGAATGTACGCTTCGAGTAGCCGTACCCCTCCTAAGATAGTAGTACTTTCAATAAAAGGAAGTCGAAATAAC

CCTGTCCCGTCGCCCCTTGCTTCCACCTTGACTCGCATGGCACACCATCATTAACTCCCGTCCATCCCATCTTCTCCTCC

CTGGATCACACGCGACGTTACACACCTGATTGAGCCCATCAGTTCGCATCGGCGCTACCAGACCTGCCAGCCCAAACC

ACGCGACGGAAACTTTGCTGCCATCGGTTGCTTGTAAAGTCGGGTATGCGGTAGATGACTGCGGCACATGGCCTCCTCC

CTCCCCCCCTTTCCGAACTTGAGTCTTGAACCGGAAGGCGTGCGTCTACAGACTTTTTCCATTAACGCGTCCGTCCGGT

GTCTGTATCTTCTCTTGACACATGTCCTTCCATACGATGTCGCCCAGTCACCATCCAACGGTGGTTTGGGGGGGGGGGG

GAGTCCCCTGCATCGTACATCGCGTGCCGCCCATCATCGCCAAAGCTTGATATCGAATTCCTGCAGCCCGGGGGATCCA

CTAGTTCTAGAGCGGCCGCCACCGCGGTGGAGCTCCAGCTTTTGTTCCCTTTAGTGAGGGTTAATTTCGAGCTTGGCGT

AATCATGGTCATAGCTGTTTCCTGTGTGAAATTGTTATCCGCTCACAATTCCACACAACATACGAGCCGGAAGCATAAA

GTGTAAAGCCTGGGGTGCCTAATGAGTGAGCTAACTCACATTAATTGCGTTGCGCTCACTGCCCGCTTTCCAGTCGGGA

AACCTGTCGTGCCAGCTGCATTAATGAATCGGCCAACGCGCGGGGAGAGGCGGTTTGCGTATTGGGCGCTCTTCCGCT

TCCTCGCTCACTGACTCGCTGCGCTCGGTCGTTCGGCTGCGGCGAGCGGTATCAGCTCACTCAAAGGCGGTAATACGG

TTATCCACAGAATCAGGGGATAACGCAGGAAAGAACATGTGAGCAAAAGGCCAGCAAAAGGCCAGGAACCGTAAAA

AGGCCGCGTTGCTGGCGTTTTTCCATAGGCTCCGCCCCCCTGACGAGCATCACAAAAATCGACGCTCAAGTCAGAGGT

GGCGAAACCCGACAGGACTATAAAGATACCAGGCGTTTCCCCCTGGAAGCTCCCTCGTGCGCTCTCCTGTTCCGACCC

TGCCGCTTACCGGATACCTGTCCGCCTTTCTCCCTTCGGGAAGCGTGGCGCTTTCTCATAGCTCACGCTGTAGGTATCTC

AGTTCGGTGTAGGTCGTTCGCTCCAAGCTGGGCTGTGTGCACGAACCCCCCGTTCAGCCCGACCGCTGCGCCTTATCC

GGTAACTATCGTCTTGAGTCCAACCCGGTAAGACACGACTTATCGCCACTGGCAGCAGCCACTGGTAACAGGATTAGC

AGAGCGAGGTATGTAGGCGGTGCTACAGAGTTCTTGAAGTGGTGGCCTAACTACGGCTACACTAGAAGGACAGTATTT

GGTATCTGCGCTCTGCTGAAGCCAGTTACCTTCGGAAAAAGAGTTGGTAGCTCTTGATCCGGCAAACAAACCACCGCT

GGTAGCGGTGGTTTTTTTGTTTGCAAGCAGCAGATTACGCGCAGAAAAAAAGGATCTCAAGAAGATCCTTTGATCTTTT

CTACGGGGTCTGACGCTCAGTGGAACGAAAACTCACGTTAAGGGATTTTGGTCATGAGATTATCAAAAAGGATCTTCA

CCTAGATCCTTTTAAATTAAAAATGAAGTTTTAAATCAATCTAAAGTATATATGAGTAAACTTGGTCTGACAGTTACCAAT

GCTTAATCAGTGAGGCACCTATCTCAGCGATCTGTCTATTTCGTTCATCCATAGTTGCCTGACTCCCCGTCGTGTAGATAA

CTACGATACGGGAGGGCTTACCATCTGGCCCCAGTGCTGCAATGATACCGCGAGACCCACGCTCACCGGCTCCAGATTT
ATCAGCAATAAACCAGCCAGCCGGAAGGGCCGAGCGCAGAAGTGGTCCTGCAACTTTATCCGCCTCCATCCAGTCTAT
TAATTGTTGCCGGGAAGCTAGAGTAAGTAGTTCGCCAGTTAATAGTTTGCGCAACGTTGTTGCCATTGCTACAGGCATC
GTGGTGTCACGCTCGTCGTTTGGTATGGCTTCATTCAGCTCCGGTTCCCAACGATCAAGGCGAGTTACATGATCCCCCAT
GTTGTGCAAAAAAGCGGTTAGCTCCTTCGGTCCTCCGATCGTTGTCAGAAGTAAGTTGGCCGCAGTGTTATCACTCATG
GTTATGGCAGCACTGCATAATTCTCTTACTGTCATGCCATCCGTAAGATGCTTTTCTGTGACTGGTGAGTACTCAACCAA
GTCATTCTGAGAATAGTGTATGCGGCGACCGAGTTGCTCTTGCCCGGCGTCAATACGGGATAATACCGCGCCACATAGC
AGAACTTTAAAAGTGCTCATCATTGGAAAACGTTCTTCGGGGCGAAAACTCTCAAGGATCTTACCGCTGTTGAGATCC
AGTTCGATGTAACCCACTCGTGCACCCAACTGATCTTCAGCATCTTTTACTTTCACCAGCGTTTCTGGGTGAGCAAAAA
CAGGAAGGCAAAATGCCGCAAAAAAGGGAATAAGGGCGACACGGAAATGTTGAATACTCATACTCTTCCTTTTTCAAT
ATTATTGAAGCATTTATCAGGGTTATTGTCTCATGAGCGGATACATATTTGAATGTATTTAGAAAAATAAACAAATAGGGG
TTCCGCGCACATTTCCCCGAAAAGTGCCAC

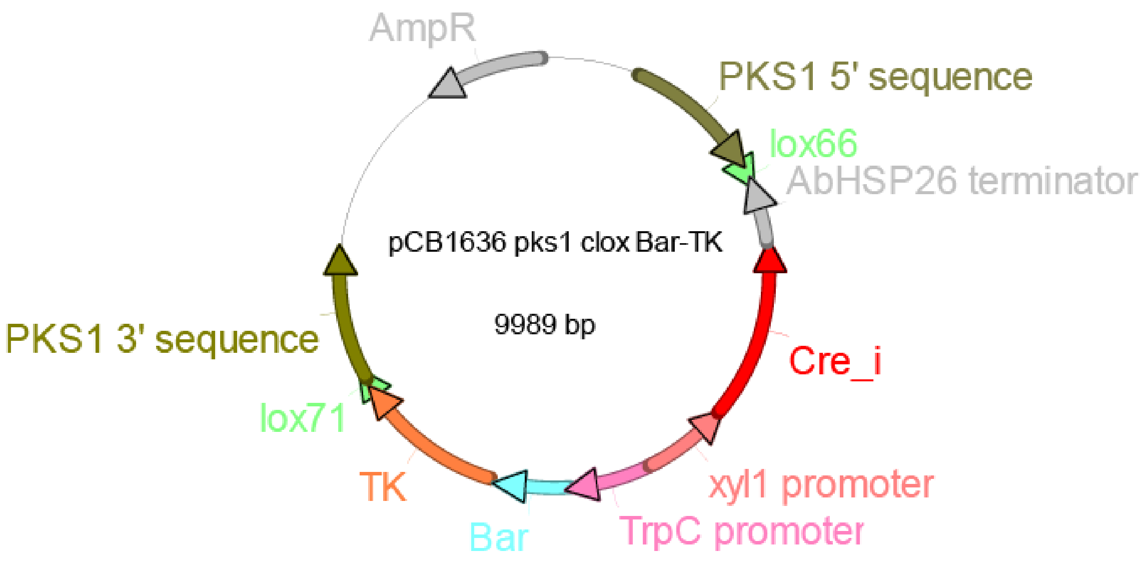


| Name | Location |
|---|---|
| PKS1 5' sequence | 675..1675 |
| lox66 | 1715..1748 |
| AbHSP26 terminator | 1760..2239 |
| Cre_i | 2240..3580 |
| xyl1 promoter | 3581..4232 |
| TrpC promoter | 4233..4917 |

| | |
|---|---|
| Bar | 4918..5466 |
| TK | 5503..6633 |
| lox71 | 6643..6676 |
| PKS1 3' sequence | 6717..7716 |
| AmpR | 9001..9861 |

> pCB1636 pks1 clox Bar-TK

CTGACGCGCCCTGTAGCGGCGCATTAAGCGCGGCGGGTGTGGTGGTTACGCGCAGCGTGACCGCTACACTTGCCAGCG
CCCTAGCGCCCGCTCCTTTCGCTTTCTTCCCTTCCTTTCTCGCCACGTTCGCCGGCTTTCCCCGTCAAGCTCTAAATCGG
GGGCTCCCTTTAGGGTTCCGATTTAGTGCTTTACGGCACCTCGACCCCAAAAAACTTGATTAGGGTGATGGTTCACGTA
GTGGGCCATCGCCCTGATAGACGGTTTTTCGCCCTTTGACGTTGGAGTCCACGTTCTTTAATAGTGGACTCTTGTTCCAA
ACTGGAACAACACTCAACCCTATCTCGGTCTATTCTTTTGATTTATAAGGGATTTTGCCGATTTCGGCCTATTGGTTAAAA
AATGAGCTGATTTAACAAAAATTTAACGCGAATTTTAACAAAATATTAACGCTTACAATTTGCCATTCGCCATTCAGGCT
GCGCAACTGTTGGGAAGGGCGATCGGTGCGGGCCTCTTCGCTATTACGCCAGCTGGCGAAAGGGGGATGTGCTGCAA
GGCGATTAAGTTGGGTAACGCCAGGGTTTTCCCAGTCACGACGTTGTAAAACGACGGCCAGTGAGCGCGCGTAATACG
ACTCACTATAGGGCGAATTGGGTACCGGGCCCCCCCTCGAGTGCCTACAAACCTGCCGTATACGGGCCATATTCGCGGG
CAGAAGGGCTCCGCCGACTCGATAATCCTGTGTGACCCTCGCGGCATCAGCGTCACGTCAACATCTAGCCTCGTTGGA
CTGGGGATGCTATGATATCAATGCATGTAGTGGACAAGCTCACGCAATGCTTACCTGCAGTTACAGTCCCTGCTGCGCG
GTGTCGAGAAGCCCGAAGCCTGGGTACGTGATACGGTACCTCGCATGCGCACCCTCGCTTCTGCTGGCTGGCGTTTTGT
TATTCTGGGCGTCTGGGATCAGAGCCAGGTAAGAAAGATGGGAAGAATGAGTGAGCACGCCTTGTGCTTTGCTCTGCC
TTGCTGCTGCGCTAGACATGCTTTGCCGTGCCGTGCTGCTGCCCTGCGCTGCGGCATGCTGCGATCGTTCCTCTTGTCTG
TCCGCCTGTCCGTCCTCCACCCCGGGCCACGATAGAGGAGCTGGCGCAAGGAAGCCCGTGTGGCGGCCTGGATCGCCT
GCCCTGAACCAATGGGTCTCGACCGATGCGGCAAGACAGCGAAAATCATCCATCAGCTTCGGTCACCGAGGAAGCTCA
GGGCCGGACTCACGCTATCCACCCGGGCAAGAAAAGGATGGGCCAAGTCAGTGGGAAAGGAAGGGGTGCAAGGGGG
GCGTGCGGATACATGGGACGTCTTCACCACACCCCCTTCTCCTTCTCTCGTTCTCACCTCACTCCCTCGACCCAGACCA
CTTGGGTTGTGTGTTCCATGTTGCAATTCCGCTATCCATCAGCCATGACTTTGACAGATGGGTTTGATTATTTTGGGGAG
ACGCCAGCTCGTGGTGGTGTTGACCTTGTCCATGGTGTGTACACCCCGTCTGTCCTTTGGCCCATCTCTTTCATATAAGT
TCCAGACTCGCTCCCAGTATCCCAATCTCAGTTTTTGTTAGGACACCGGGAAACCTCTCAAACACCTAGTAGGAAATTC
ATGTCGACAACCTGACAAGGTCGACGTTAACTGATATTGAAGGAGCATTTTTTGGGCTTACCGTTCGTATAGCATACATT
ATACGAAGTTATTGGGGCGCGCCTTGAACAGCATTGCGTTATTTATAACTTAGAAGAGAATCACAAGGCATGTTAAGGC
TTAAGCCGTGGAGCGTACTATTTGGGGTGTCTAGTACGCGAAGGTAAATTTCAGATGTGCTTCGTGCGGTACAGTGTGG
TCTATCCGGAACAGCCCTGTACTGCGGAACTTGAGTCGTACAGCCATTTCACTTTGAGTGCGGAAGGACCATTTAGGAG
GTAAATTTGGTAATGGTAAATGCAGATTAAGAAAGGTATTATTTACGATTACAAGATGCTAGAGAAAATAAAAGAATACT
CTGAAGTAGGGAGTCGAGTCCGATAAAATACATCATAATGGATACAACATGTGCAAGAGGGGAGGTGAAAGTGTTTCT
ATCAAAAAATAATCATATAATGACGACCAAGAAAGCTAAACTCGCAATGCATAAAACAAGGCCATTTAAAACAATCGAA
ATAAAAGTCCAAAACCGAGGAGCTACTAAGAGCTCTTAATCGCCGTCTTCGAGGAGACGGACCATAGCGCCGGTCTCG

CTGTCGAGGTTGCGGATGTAGTTCATGACGATGTTGACGTTGGTCCAGCCTCCGGCTTGCATGATCTCAGGAATGGAGA
CGCCGGCGCGGGCCATGTCGCGGGCGGCGCCCACGCGGGCGGAGTGGCCGGACCAGGCGAGGTAGCGCTGGCCAGA
ATCGTCCTTGGCACCGTAGATCAGGCGATGGGTGGCTTCGAAGATACCCTCCAGGGCGCGGGTCGAGAGTTGGGACGT
GGCGCTAGGGGCAGCGACGCCGTTCTTGCGCACGCGACAGAAGAGGTAGTTGTTAGGGTCGTCGGCAACGCCCGAAA
CGCTAATCCAGCGTTCGACGAGCTTAGTGACACCGAGGCTGAGCGCCTTTTCCACACCAGCTGTGCTCACCAAAGTTT
TGGTTCTTCCGATGTGAATGAGCATGCCTGAAAAAGATCAGTACGATGTTTGATAGAATGAAATAGGAAGTCGTGGTGA
AGCCCGGGTTAAGCCTGGGAAGACGCGTCTAGATGCGTGATGCATGCCAACGGGCAAGTGCCCATTACACGGCGGGGT
ACGCCACGATGCGAGATGCGAGACGCGCGCCAAAACCGACAGCGACGACGAGGTATAATAAAAATGGTACTTACGTCC
GCCATCGGTTCGGCTGATGTCCTTCACGCGAATGCGCGCGATTTCGGCGATGCGGAGCAAGGTGTTGTACGCGATACCC
AGAAAAGCAAGATTTCGGATGTCCTGGCAGCGGTCCGAGTTCTCCATAAGAGATCGGACCTGGTCGAAATCCGTTCTC
TCGAAAGCCAGAGCTTGCTTAGCTCGTTCTCCCGCATCGACGTTTTCTTTTCGGATTCGTCTCATAACCAGCGAGACCG
CATTCGAGTCGCTGGGTCGGGGAAGACCCGAACGTCTGTGAAGCATGTTCAACTGTCCCAAGTGCTGCTGAATGGTCT
TGACAGCCAGGCCTCTAGCCTGCAAATACAGCAGATAGTCACGGACGTCCTCTGGCTCGGCAGGAAACCACTTTCGGT
TGTTCAACTTGCACCATGCTGCCCAGGAACGGCAGACGCTCAGGAGCATCTTCCAAGTATGCTCGGAAAAGGCTTGGC
GATCTCGGAACATGTCCATCAAGTTCTTTCGGACTTCATCGGAGGTTGCATCAACCGGCAGGGCCGGGAGATTCTGGTG
GACTGTCAACAGGTTGCTCTCAAGCAGCTCCTTCAGGTGCGCCGCCTTTCGATAACGGCTACGTCTGGAGGACTCTCTA
TTCGACTCCTTTCTCTTACGAACGCGCTCTTCCGTTGCCATGGGATTGTAAGGGGACGATGATGAAGTTGATAGCATTCA
AAGATGTTGCTCGTACAAGTGGACTGTCGACATCTCCGGGTTTAATATTGCTCTCTACTGCTACTCACTGTCAAGGTATC
TGGCTAAAATGCAGGTCAGTCGTGACCAAACCTCCATTGCTCCGAGAAACCTCTGATGAGGAAAACGAAATTCACCAT
TGTTGCATAACATAGTGTTACACCCAGTCGGGTATGGCATGCCGCGCGCCAGCGATGAGAGATGCATCGAGTGTTGTGA
TTACTTGGCTTCGTGACATGTGAGGCCGGCTAAATAGGCCCCTTGAGTACACTAGACCAACTTGAACTGGGGTCATCGG
TAGTTGGCTGTCACGGCATCTGCTCGTAAAGACCTGCATCCCGTCCCGTTCTTGCAGCGGTTAAATGCGATCGAGTTAC
TTTTATCCTGCCAGGGTACCGGTGCTGGGCAAGGTGTACGCAAGGTTGAACACTCAGGAATAGATACCAAGCAAGAAT
GAGGGCATCGTCAGCCGAGATGGCGACTGCAACAGCACTTGTCCCTAGTGGTAGCTCCTGGAGAACTGATCATTGTGT
GAAGCATTATGATCCACATTCAGGCTGTTTAATACCCTGCTAAGGGTAAATCCGGCAAGTTTGCTAGCGAGCTAGTGGA
GGTCAACAATGAATGCCTATTTTGGTTTAGTCGTCCAGGCGGTGAGCACAAAATTTGTGTCGTTTGACAAGATGGTTCA
TTTAGGCAACTGGTCAGATCAGCCCCACTTGTAGCAGTAGCGGCGGCGCTCGAAGTGTGACTCTTATTAGCAGACAGG
AACGAGGACATTATTATCATCTGCTGCTTGGTGCACGATAACTTGGTGCGTTTGTCAAGCAAGGTAAGTGGACGACCCG
GTCATACCTTCTTAAGTTCGCCCTTCCTCCCTTTATTTCAGATTCAATCTGACTTACCTATTCTACCCAAGCATCCAAATG
AAAAAGCCTGAACTCACCGCGACGTCTGTCGAGAAGTTTCTGATCGAAAAGTTCGACAGCGTCTCCGACCTGATGCA
GCTCTCGGAGGGCGAAGAATCTCGTGCTTTCAGCTTCGATGTAGGAGGGCGTGGATATGTCCTGCGGGTAAATAGCTGC
GCCGATGGTTTCTACAAAGATCGTTATGTTTATCGGCACTTTGCATCGGCCGCGCTCCCGATTCCGGAAGTGCTTGACAT
TGGGGAGTTCAGCGAGAGCCTGACCTATTGCATCTCCCGCCGTGCACAGGGTGTCACGTTGCAAGACCTGCCTGAAAC
CGAACTGCCCGCTGTTCTCCAGCCGGTCGCGGAGGCCATGAGCCCAGAACGACGCCCGGCCGACATCCGCCGTGCCA
CCGAGGCGGACATGCCGGCGGTCTGCACCATCGTCAACCACTACATCGAGACAAGCACGGTCAACTTCCGTACCGAGC
CGCAGGAACCGCAGGAGTGGACGGACGACCTCGTCCGTCTGCGGGAGCGCTATCCCTGGCTCGTCGCCGAGGTGGAC

GGCGAGGTCGCCGGCATCGCCTACGCGGGTCCCTGGAAGGCACGCAACGCCTACGACTGGACGGCCGAATCGACCGT
GTACGTCTCCCCCCGCCACCAGCGGACGGGACTGGGCTCCACGCTCTACACCCACCTGCTGAAGTCCCTGGAGGCACA
GGGCTTCAAGAGCGTGGTCGCTGTCATCGGGCTGCCCAACGACCCGAGCGTGCGCATGCACGAGGCGCTCGGATATGC
CCCCCGCGGCATGCTGCGGGCGGCCGGCTTCAAGCACGGGAACTGGCATGACGTGGGTTTCTGGCAGCTGGACTTCA
GCCTGCCGGTTCCGCCCCGTCCGGTCCTGCCCGTCACCGAGATTATCGATGGAGCTGGAGCAGGAGCTGGAGCCGGTG
CGATGGCTTCCTACCCCTGCCATCAGCACGCCTCCGCCTTTGACCAGGCGGCTAGATCCCGCGGACATAGCAACCGCA
GAACAGCCCTCCGACCGCGCCGTCAACAGGAAGCGACGGAAGTTCGATTGGAGCAGAAGATGCCCACCCTCCTTAGA
GTCTATATCGACGGCCCTCACGGAATGGGCAAGACGACCACCACCCAGCTCCTCGTCGCACTGGGTAGCCGTGATGAC
ATCGTCTACGTTCCTGAGCCAATGACGTATTGGCAAGTGCTCGGCGCATCCGAAACCATTGCCAACATTTACACCACCC
AACACCGCCTGGACCAGGGTGAAATTTCCGCGGGAGATGCTGCCGTCGTCATGACTTCGGCCCAAATTACTATGGGTAT
GCCGTACGCCGTCACGGACGCTGTCCTTGCACCCCACGTGGGCGGAGAGGCCGGTTCGTCTCATGCCCCTCCTCCTGC
CCTCACCCTTATCTTCGATCGCCACCCTATTGCCGCGTTGCTCTGTTACCCTGCTGCCCGCTACCTCATGGGTTCTATGAC
ACCGCAGGCTGTCCTTGCTTTCGTCGCTCTCATCCCGCCAACACTGCCCGGCACTAATATCGTCCTTGGCGCTCTCCCA
GAAGACCGACACATTGATAGACTCGCCAAGCGCCAGAGACCTGGAGAGAGATTGGACTTGGCTATGCTCGCCGCCATC
CGTCGTGTCTACGGCCTCCTCGCTAACACAGTCAGATATCTGCAGGGAGGCGGATCCTGGTGGGAGGACTGGGGTCAG
CTTTCGGGTACTGCTGTGCCTCCCCAGGGCGCCGAGCCGCAGAGCAACGCCGGTCCCAGACCCCATATCGGTGACACG
CTCTTTACCCTGTTCCGCGCGCCTGAGCTGCTGGCCCCGAACGGAGATTTGTACAACGTCTTCGCCTGGGCCTTGGACG
TGCTCGCTAAGCGACTCCGCCCAATGCACGTGTTCATCTTGGACTACGACCAATCTCCTGCCGGTTGCCGCGACGCCCT
CCTGCAATTGACCTCTGGCATGGTTCAGACCCACGTCACTACACCGGGAAGCATCCCTACGATCTGCGATCTTGCCCGC
ACATTCGCACGAGAGATGGGCGAAGCGAATTAAAGTCATATGATAACTTCGTATAGCATACATTATACGAACGGTAAGAT
GCCGACCGGGAACCAGTTAACGTCGACGGTATCGATGCCAAGCAACTTGGAACTTTCCTTCGTGAAGCGGTTTGCGAG
TAAGGGATTCACATAGCGAGGTTGCATACAAGCACGCGCATAGCAGGTATAGAAGGGAAACAGGACATTACATGATTG
CGAGCATGGATATGAGACTGCCTCTTTTTTTCCATCATTGTTCATTTTTCTTGTCATCTTTTCATCTTTCTTTTTTACCACC
CTGTCTTAACATTGATATCCGCTCCCTGTACAAGCGCGAGATCATGCACACTCTTACGAATCATTCCTTACTCAACGTCG
GCCGGAAAGGGAGGTAGAGGTGGAATTATCTGGGGAAACAGCATGGTCAACACCTCAGCCTTCGACCTTCTTGGGCA
CGGGAAGGACGGATGGCGCGGGTGTTGGCTTCGAAAGTAATTGGGCTCCATCATTTTCTTTTGTCACGAATTAGCAGAT
TCCTCATGTCAGGCGGTGTTTACCAGGTTTTGTCGTCTTTTTCTGCGGTCCATAGAATGTACGCTTCGAGTAGCCGTACC
CCTCCTAAGATAGTAGTACTTTCAATAAAAGGAAGTCGAAATAACCCTGTCCCGTCGCCCCTTGCTTCCACCTTGACTC
GCATGGCACACCATCATTAACTCCCGTCCATCCCATCTTCTCCTCCCTGGATCACACGCGACGTTACACACCTGATTGAG
CCCATCAGTTCGCATCGGCGCTACCAGACCTGCCAGCCCAAACCACGCGACGGAAACTTTGCTGCCATCGGTTGCTTG
TAAAGTCGGGTATGCGGTAGATGACTGCGGCACATGGCCTCCTCCCTCCCCCCCTTTCCGAACTTGAGTCTTGAACCGG
AAGGCGTGCGTCTACAGACTTTTTCCATTAACGCGTCCGTCCGGTGTCTGTATCTTCTCTTGACACATGTCCTTCCATAC
GATGTCGCCCAGTCACCATCCAACGGTGGTTTGGGGGGGGGGGGGAGTCCCCTGCATCGTACATCGCGTGCCGCCCAT
CATCGCCAAAGCTTGATATCGAATTCCTGCAGCCCGGGGGATCCACTAGTTCTAGAGCGGCCGCCACCGCGGTGGAGC
TCCAGCTTTTGTTCCCTTTAGTGAGGGTTAATTTCGAGCTTGGCGTAATCATGGTCATAGCTGTTTCCTGTGTGAAATTG
TTATCCGCTCACAATTCCACACAACATACGAGCCGGAAGCATAAAGTGTAAAGCCTGGGGTGCCTAATGAGTGAGCTA

ACTCACATTAATTGCGTTGCGCTCACTGCCCGCTTTCCAGTCGGGAAACCTGTCGTGCCAGCTGCATTAATGAATCGGC

CAACGCGCGGGGAGAGGCGGTTTGCGTATTGGGCGCTCTTCCGCTTCCTCGCTCACTGACTCGCTGCGCTCGGTCGTT

CGGCTGCGGCGAGCGGTATCAGCTCACTCAAAGGCGGTAATACGGTTATCCACAGAATCAGGGGATAACGCAGGAAAG

AACATGTGAGCAAAAGGCCAGCAAAAGGCCAGGAACCGTAAAAAGGCCGCGTTGCTGGCGTTTTTCCATAGGCTCCG

CCCCCCTGACGAGCATCACAAAAATCGACGCTCAAGTCAGAGGTGGCGAAACCCGACAGGACTATAAAGATACCAGG

CGTTTCCCCCTGGAAGCTCCCTCGTGCGCTCTCCTGTTCCGACCCTGCCGCTTACCGGATACCTGTCCGCCTTTCTCCCT

TCGGGAAGCGTGGCGCTTTCTCATAGCTCACGCTGTAGGTATCTCAGTTCGGTGTAGGTCGTTCGCTCCAAGCTGGGCT

GTGTGCACGAACCCCCCGTTCAGCCCGACCGCTGCGCCTTATCCGGTAACTATCGTCTTGAGTCCAACCCGGTAAGACA

CGACTTATCGCCACTGGCAGCAGCCACTGGTAACAGGATTAGCAGAGCGAGGTATGTAGGCGGTGCTACAGAGTTCTT

GAAGTGGTGGCCTAACTACGGCTACACTAGAAGGACAGTATTTGGTATCTGCGCTCTGCTGAAGCCAGTTACCTTCGGA

AAAAGAGTTGGTAGCTCTTGATCCGGCAAACAAACCACCGCTGGTAGCGGTGGTTTTTTTGTTTGCAAGCAGCAGATT

ACGCGCAGAAAAAAAGGATCTCAAGAAGATCCTTTGATCTTTTCTACGGGGTCTGACGCTCAGTGGAACGAAAACTCA

CGTTAAGGGATTTTGGTCATGAGATTATCAAAAAGGATCTTCACCTAGATCCTTTTAAATTAAAAATGAAGTTTTAAATC

AATCTAAAGTATATATGAGTAAACTTGGTCTGACAGTTACCAATGCTTAATCAGTGAGGCACCTATCTCAGCGATCTGTC

TATTTCGTTCATCCATAGTTGCCTGACTCCCCGTCGTGTAGATAACTACGATACGGGAGGGCTTACCATCTGGCCCCAGT

GCTGCAATGATACCGCGAGACCCACGCTCACCGGCTCCAGATTTATCAGCAATAAACCAGCCAGCCGGAAGGGCCGAG

CGCAGAAGTGGTCCTGCAACTTTATCCGCCTCCATCCAGTCTATTAATTGTTGCCGGGAAGCTAGAGTAAGTAGTTCGC

CAGTTAATAGTTTGCGCAACGTTGTTGCCATTGCTACAGGCATCGTGGTGTCACGCTCGTCGTTTGGTATGGCTTCATTC

AGCTCCGGTTCCCAACGATCAAGGCGAGTTACATGATCCCCCATGTTGTGCAAAAAAGCGGTTAGCTCCTTCGGTCCTC

CGATCGTTGTCAGAAGTAAGTTGGCCGCAGTGTTATCACTCATGGTTATGGCAGCACTGCATAATTCTCTTACTGTCATG

CCATCCGTAAGATGCTTTTCTGTGACTGGTGAGTACTCAACCAAGTCATTCTGAGAATAGTGTATGCGGCGACCGAGTT

GCTCTTGCCCGGCGTCAATACGGGATAATACCGCGCCACATAGCAGAACTTTAAAAGTGCTCATCATTGGAAAACGTTC

TTCGGGGCGAAAACTCTCAAGGATCTTACCGCTGTTGAGATCCAGTTCGATGTAACCCACTCGTGCACCCAACTGATCT

TCAGCATCTTTTACTTTCACCAGCGTTTCTGGGTGAGCAAAAACAGGAAGGCAAAATGCCGCAAAAAAGGGAATAAG

GGCGACACGGAAATGTTGAATACTCATACTCTTCCTTTTTCAATATTATTGAAGCATTTATCAGGGTTATTGTCTCATGAG

CGGATACATATTTGAATGTATTTAGAAAAATAAACAAATAGGGGTTCCGCGCACATTTCCCCGAAAAGTGCCAC

Figure S2. The vector information used in this study.
